# High and stable ATP levels prevent aberrant intracellular protein aggregation

**DOI:** 10.1101/801738

**Authors:** Masak Takaine, Hiromi Imamura, Satoshi Yoshida

**Author notes:** Correspondence should be addressed to (M.T.) (Lead contact) or (S.Y.).

## Abstract

ATP at millimolar levels has recently been implicated in the solubilization of cellular proteins. However, the significance of this high ATP level under physiological conditions and the mechanisms that maintain ATP remain unclear. We herein demonstrated that AMP-activated protein kinase (AMPK) and adenylate kinase (ADK) cooperated to maintain cellular ATP levels regardless of glucose levels. Single cell imaging of ATP-reduced yeast mutants revealed that ATP levels in these mutants repeatedly underwent stochastic and transient depletion, which induced the cytotoxic aggregation of endogenous proteins and pathogenic proteins, such as huntingtin and α-synuclein. Moreover, pharmacological elevations in ATP levels in an ATP-reduced mutant prevented the accumulation of α-synuclein aggregates and its cytotoxicity. The removal of cytotoxic aggregates depended on proteasomes, and proteasomal activity cooperated with AMPK or ADK to resist proteotoxic stresses. The present study is the first to demonstrate that cellular ATP homeostasis ensures proteostasis and revealed that suppressing the high volatility of cellular ATP levels prevented cytotoxic protein aggregation, implying that AMPK and ADK are important factors that prevent proteinopathies, such as neurodegenerative diseases.

## INTRODUCTION

Adenosine triphosphate (ATP) is the main energy currency used by all living organisms. In addition to its role as energy currency, ATP has recently been suggested to influence the balance between the soluble and aggregated states of proteins, indicating that proteostasis is maintained by energy-dependent chaperones and also by the property of ATP as a hydrotrope to solubilize proteins (Hayes, Peuchen et al., 2018, Patel, Malinovska et al., 2017, Pu, Li et al., 2019, Sridharan, Kurzawa et al., 2019). Furthermore, ATP levels have been shown to regulate the physicochemical properties of the cytoplasm, such as viscosity, macromolecular crowding, and liquid-liquid phase separation (Marini, Nuske et al., 2020, Persson, Ambati et al., 2020). However, the role of ATP was assessed in these studies using artificial ATP depletion. Therefore, it currently remains unclear whether ATP-dependent protein solubilization/desolubilization have physiologically significant cellular roles.

We recently established a reliable imaging technique to quantify intracellular ATP levels in single living yeast cells using the genetically encoded fluorescent ATP biosensor QUEEN (Yaginuma, Kawai et al., 2014), which enables long-term evaluations of ATP homeostasis in individual cells (Takaine, Ueno et al., 2019). The findings obtained demonstrated that intracellular ATP levels did not vary within a yeast population grown in the same culture (Takaine et al., 2019), which was in contrast to the large variations observed in intracellular ATP levels within a bacterial cell population (Yaginuma et al., 2014). Moreover, intracellular ATP levels in individual living yeast cells were stably and robustly maintained at approximately 4 mM, irrespective of carbon sources and cell cycle stages, and temporal fluctuations in intracellular ATP levels were small (Takaine et al., 2019). Based on these findings, we hypothesized that an exceptionally robust mechanism exists to precisely regulate ATP levels in eukaryotes. It currently remains unclear why ATP is stably maintained at a markedly higher level than the *K*_*m*_ (Michaelis constant) required for the enzymatic activities of most ATPases (Edelman, Blumenthal et al., 1987), and the consequences associated with failed ATP homeostasis in living organisms have not yet been elucidated.

The most promising candidate regulator of ATP homeostasis is AMP-activated protein kinase (AMPK). AMPK, which is activated by AMP and inhibited by ATP (Xiao, Heath et al., 2007), has long been regarded as an important regulator of the whole-body and cellular energy status in eukaryotes (Hardie, Schaffer et al., 2016). AMPK is activated by increases in the AMP:ATP or ADP:ATP ratio (*i*.*e*., low-energy state), and regulates its downstream effectors by phosphorylation to redirect cell metabolism from an anabolic (ATP-consuming) state to catabolic (ATP-producing) state (Herzig & Shaw, 2017). In the budding yeast *Saccharomyces cerevisiae*, the sucrose non-fermenting 1 (Snf1) protein kinase complex is the sole AMPK. Similar to other AMPKs, the yeast Snf1 complex comprises three subunits: the catalytic α subunit (*SNF1*), scaffolding β subunit (*SIP1, SIP2* or *GAL83*), and regulatory γ subunit (*SNF4*) (Ghillebert, Swinnen et al., 2011). The role of the Snf1 complex in adaptation to glucose limitations has been characterized in detail (Hedbacker & Carlson, 2008). The Snf1 complex is inactive in the presence of sufficient glucose levels in media (Wilson, Hawley et al., 1996). Decreases in glucose levels have been shown to activate the Snf1 complex and phosphorylate the transcriptional repressor Mig1, which then triggers the transcription of numerous glucose-repressed genes (Carlson, 1999). However, the contribution of AMPK or the Snf1 complex to cellular ATP levels remains unknown.

Other possible candidate regulators of ATP homeostasis include genes whose mutation leads to decreases in the cellular content of ATP. However, based on biochemical analyses of cell populations, few yeast mutants reduced ATP levels (Gauthier, Coulpier et al., 2008, Ljungdahl & Daignan-Fornier, 2012). Adenylate kinase (ADK) is a key enzyme that synthesizes ATP and AMP using two ADP molecules as substrates, and the null mutant of ADK (*adk1Δ*) was shown to have a reduced cellular ATP level (∼70% of the wild type) (Gauthier et al., 2008). Bas1 is a transcription factor that is required for *de novo* purine synthesis and *bas1Δ* also has a reduced ATP level (∼50% of the wild type) (Gauthier et al., 2008). However, the regulation of ATP levels and the physiological consequences of reduced ATP levels in these mutants remain unclear, particularly at the single cell level.

In the present study, we investigated the roles of AMPK, ADK, and Bas1 in ATP homeostasis using the QUEEN-based single cell ATP imaging technique. We demonstrated for the first time that AMPK is involved in the regulation of intracellular ATP levels, even under glucose-rich conditions. Furthermore, time-lapse ATP imaging revealed that cells lacking both AMPK and ADK frequently underwent transient ATP depletion, while ATP levels oscillated in those lacking Bas1. These ATP dynamics in the mutants were overlooked in previous biochemical studies. We found that some intrinsic proteins and aggregation-prone model proteins, including α-synuclein, which is responsible for Parkinson’s disease, aggregated and were cytotoxic in all of the ATP-reduced mutants tested. The present results suggest that the stable maintenance of ATP is essential for proteostasis and imply that an ATP crisis promotes proteinopathies, such as neurodegenerative diseases.

## RESULTS

### ADK1 cooperates with AMPK to regulate ATP homeostasis

We recently developed a reliable monitoring system for cytoplasmic ATP levels in living yeast cells using the ATP biosensor QUEEN (Takaine et al., 2019). We herein conducted a more detailed examination of ATP dynamics in wild-type and mutant yeast cells using this system. We initially investigated whether the deletion of *SNF1*, which encodes a catalytic subunit of AMPK, affected cellular ATP levels. Cellular ATP levels were significantly lower in *SNF1*-null mutant (*snf1Δ*) cells than in wild-type cells at various glucose levels (Fig. 1A and Fig. S1A, B). It is important to note that in addition to low glucose conditions (0.05% glucose), at which the Snf1 complex is active, *snf1*Δ cells showed significantly reduced ATP levels even under high glucose conditions (2% glucose), at which the Snf1 complex is considered to be inactive. The deletion of *MIG1* had a negligible effect on ATP levels (Fig. S1C), suggesting that as yet unknown factors other than Mig1 primarily regulate ATP levels under the control of the Snf1 complex. Collectively, these results demonstrated for the first time that AMPK/SNF1 affect cellular ATP levels even under glucose-rich conditions.

**Fig 1.**
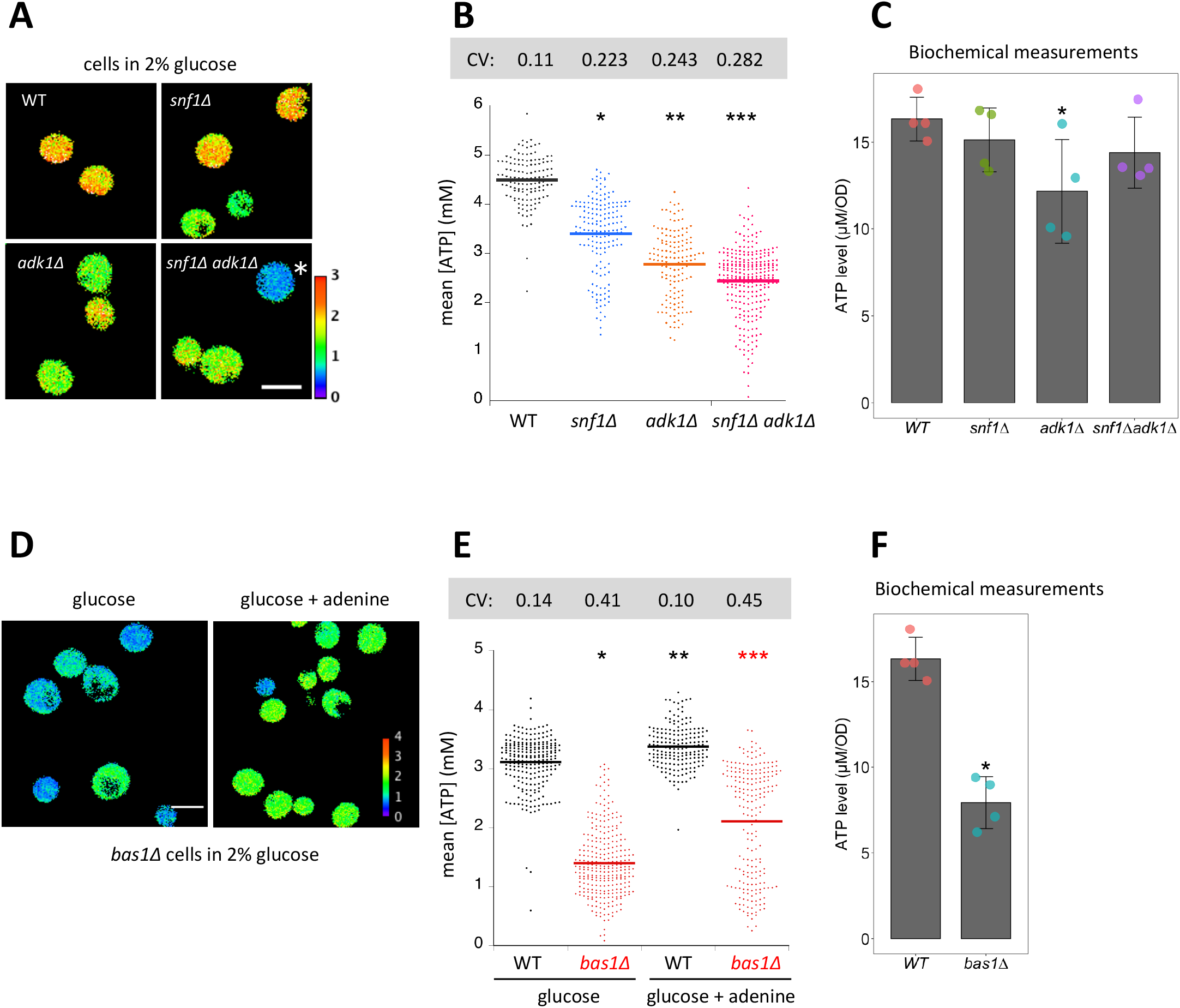
Interconversion and active synthesis of adenine nucleotides are important for ATP homeostasis. (A) Adk1 and Snf1 synergistically control cellular ATP levels. QUEEN ratio images of ATP homeostasis mutant cells grown in medium containing 2% glucose. The asterisk indicates an example of cells with significantly reduced ATP levels. (B) The mean QUEEN ratios of cells were translated to ATP levels and shown in a dot plot. The horizontal bar indicates the mean of each population. Asterisks indicate *p* values versus WT: *=1.5×10^−59^, **=2.3×10^−34^, ***=2.9×10^−117^. CV: coefficient of variance. N = 134–276 cells were scored. (C) Biochemical measurements of cellular ATP levels. ATP levels in cells of the indicated genotypes were measured as described in the Materials and Methods. Data are the mean ± 1SD (error bars) (N = 4). An asterisk indicates a *p* value of 0.03 versus WT. (D) QUEEN ratio images of *bas1Δ* cells grown in 2% glucose medium. Growth in media supplemented with 0.11 mg/ml adenine partially restored the low ATP phenotype of *bas1*Δ. (E) ATP levels in cells shown in *D* were plotted. Asterisks indicate *p* values: *=2.3×10^−160^, **=8.8×10^−12^ (versus WT in glucose), ***=3.6×10^−20^ (versus *bas1*Δ in glucose). CV: coefficient of variance. N = 186–296 cells were scored. (F) ATP levels in WT and *bas1Δ* cells were measured as described in *C*. Data are the mean ± 1SD (error bars) (N = 4). An asterisk indicates a *p* value of 8.5×10^−5^ versus WT.

ADK catalyzes the interconversion of adenine nucleotides (ATP+AMP ⟷ 2ADP), which is important for *de novo* adenine nucleotide synthesis and the balance between ATP, ADP, and AMP. Previous biochemical studies reported that the deletion of the ADK gene reduced ATP levels in mouse skeletal muscle cells and budding yeasts (Gauthier et al., 2008, Janssen, Dzeja et al., 2000). We confirmed these findings using an ATP imaging method: *adk1*Δ cells showed significantly lower QUEEN ratios than wild-type cells on average in the presence of sufficient carbon sources (Fig. S2).

In addition to being a key enzyme in purine metabolism, ADK has also been suggested to cooperate with AMPK in order to monitor the cellular energy state (Hardie, Carling et al., 1998). Therefore, we compared ATP levels in *snf1*Δ *adk1*Δ double mutant cells with those in *snf1*Δ, *adk1*Δ cells and wild-type cells (Fig. 1A, B). *snf1Δ adk1Δ* cells had significantly lower average ATP levels than single mutant cells. We also found not only a general reduction, but also a marked variation in ATP levels in the *snf1*Δ, *adk1Δ, snf1*Δ *adk1Δ* cell population, as indicated by the large coefficient of variance (CV) (Fig. 1B). Furthermore, some *snf1Δ adk1Δ* cells had very low ATP levels (Fig. 1A, B). These results suggest that Adk1 and the Snf1 complex both synergistically contribute to ATP homeostasis.

We confirmed the decreases observed in ATP levels in *snf1Δ, adk1Δ* and *snf1Δadk1Δ* cells using a biochemical assay of whole cell extract (Fig. 1C). ADP levels were also reduced in these mutant cells (Fig. S3). In *adk1Δ* cells, the ATP/ADP ratio increased, whereas the sum of ATP and ADP levels decreased (Fig. S3), which is consistent with previous findings (Gauthier et al., 2008). These biochemical data and their relevance to QUEEN data are discussed later.

### A large pool of adenine nucleotides is important for maintaining cellular ATP levels

We examined a *bas1Δ* mutant, which is defective in the expression of genes responsible for adenine biogenesis (Daignan-Fornier & Fink, 1992, Denis, Boucherie et al., 1998). Consistent with previous biochemical findings (Gauthier et al., 2008), ATP levels quantified by QUEEN were reduced by ∼50% in *bas1Δ* cells (Fig. 1D, E). We found not only a general reduction, but also a marked variation in ATP levels in the *bas1Δ* cell population, as indicated by the large CV (Fig. 1E). The decrease observed in ATP levels was due to reduced adenine biosynthesis because the addition of extra adenine to media partially restored ATP levels (Fig. 1E). These results suggest that the sufficient production of adenine nucleotides is essential for the stable maintenance of ATP levels. Moreover, the role of Bas1 in maintaining ATP levels appeared to be epistatic to that of Snf1 because *bas1Δ snf1Δ* double mutant cells showed a similar distribution of ATP levels to *bas1Δ* cells (Fig. S4). We confirmed the decrease observed in ATP levels in *bas1Δ* cells using a biochemical assay (Fig. 1F). ADP levels and the sum of ATP and ADP levels also significantly decreased in *bas1Δ* cells (Fig. S3), which is consistent with previous findings (Gauthier et al., 2008).

### ATP levels temporally fluctuate in ATP mutant cells

To investigate the mechanisms contributing to the marked variations in ATP levels in *snf1Δ adk1Δ* cells in more detail, we employed time-lapse ATP imaging (Fig. 2). We found that the QUEEN ratio often underwent a rapid decline followed by recovery in *snf1Δ adk1Δ* cells (see 116 and 132 min in Fig. 2A, C, and Movie S1, and 180 and 356 min in Fig. 2B, D, and Movie S2). The sudden decrease in ATP levels (hereafter called “the ATP catastrophe”) occurred within a few minutes without any sign and was never observed in wild-type cells (Takaine et al., 2019). The ATP catastrophe appeared to be a stochastic event and cell intrinsic: these events occurred independent of the cell cycle stage or cell size (compare Fig. 2C with D). Under some conditions, the QUEEN ratio did not recover after the ATP catastrophe and the cell died, as judged by the loss of QUEEN signals in the cell (Fig. S5). These results suggest that the marked variations observed in ATP levels in *snf1Δ adk1Δ* cells were not simply due to a mixed population with different basal ATP levels, but were rather caused by the stochastic ATP catastrophe in individual cells.

**Fig 2.**
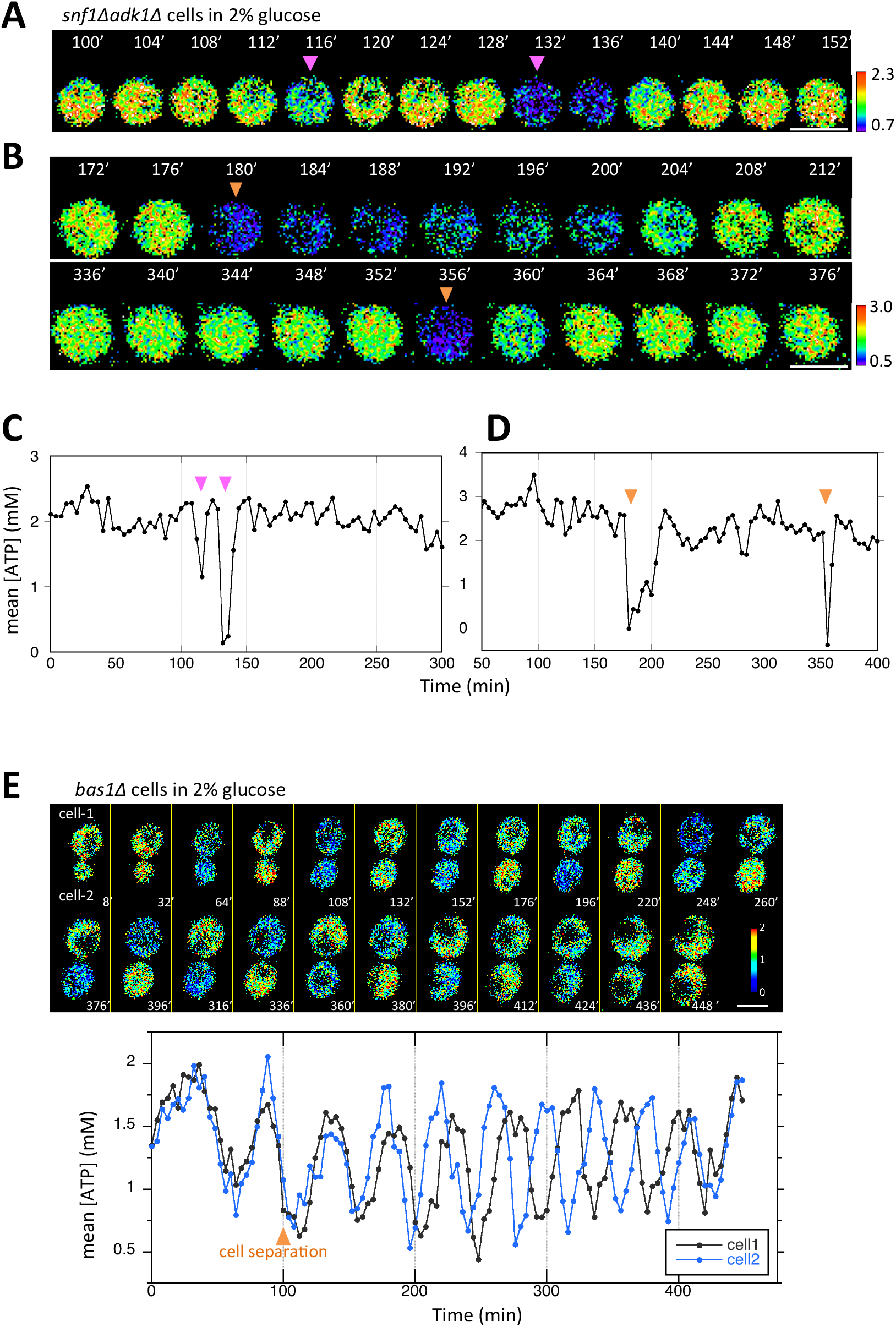
Temporal fluctuations in ATP levels in *snf1Δadk1Δ* and *bas1Δ* cells. (A, C) Time-lapse imaging of QUEEN in *snf1*Δ *adk1*Δ cells in 2% glucose medium. Images at the representative time points were shown. The QUEEN ratio decreased twice (indicated by arrowheads) within a short interval. See also Movie S1. Data were converted into ATP levels and plotted in (C). (B, D) Another example of the time-lapse imaging of QUEEN in *snf1*Δ*adk1*Δ cells in 2% glucose medium. The QUEEN ratio decreased twice (indicated by arrowheads) with a long interval. See also Movie S2. Data were converted into ATP levels and plotted in (D). (E) Time-lapse imaging of QUEEN in *bas1*Δ in 2% glucose medium. ATP levels in the mother (cell-1) and daughter (cell-2) were plotted at the bottom. Images at the representative time points were shown on the top. Note that the QUEEN ratio is synchronized until cells undergo separation at the time point of 76 min indicated by an arrow. After separation, each cell has a unique periodic cycle of ATP. The movie is available in Movie S3. White scale bar = 5 µm.

Time-lapse imaging of *bas1Δ* revealed oscillatory cycles in ATP levels (Figs. 2E and S6A, and Movies S3 and S4): ATP cycling in *bas1Δ* cells was slow (∼35 min on average, Fig. S6B) and distinguishable from that in *snf1Δ adk1Δ* cells; however, the common characteristics of these mutants were that the level of ATP repeatedly reached close to 0 mM. The ATP oscillation cycle was unsynchronized in the population and independent of cell cycle progression, suggesting a unique metabolic rhythm intrinsic to each cell. The oscillatory nature of ATP cycling in the *bas1Δ* mutant may involve a transcription/translation cycle and will be described elsewhere.

### ATP homeostasis is required for preventing protein aggregation *in vivo*

We recently reported that cellular ATP levels were stably maintained at ∼4 mM in budding yeast (Takaine et al., 2019) and herein demonstrated that Adk1 and Bas1 in the Snf1/AMPK complex were required for the regulation of ATP homeostasis. However, the physiological importance of ATP homeostasis remains unknown. To clarify the significance of high ATP levels, we examined the global genetic interactions of *snf1Δ, adk1Δ*, and *bas1Δ* using CellMap ((Usaj, Tan et al., 2017), thecellmap.org). An *in silico* analysis identified genes involved in “protein folding/glycosylation” as common negative genetic interactors with *adk1Δ* and *bas1Δ* (Fig. 3A). Negative genetic interactors of *ura6*, a gene encoding uridylate kinase that also exhibits ADK activity, were enriched in the “protein folding/glycosylation” category (Fig. 3A). We also found that interactors of *snf1* were implicated in “protein folding/glycosylation”. None of these mutants exhibited apparent genetic interactions with genes in the “metabolism” category (Fig. 3A). The same analysis using genetic and physical interactors provided similar results and showed that many interactors were enriched in the “protein turnover” category (Fig. S7). These results imply that although these three mutants regulate ATP with distinct mechanisms, all three have a common cellular function.

**Fig 3.**
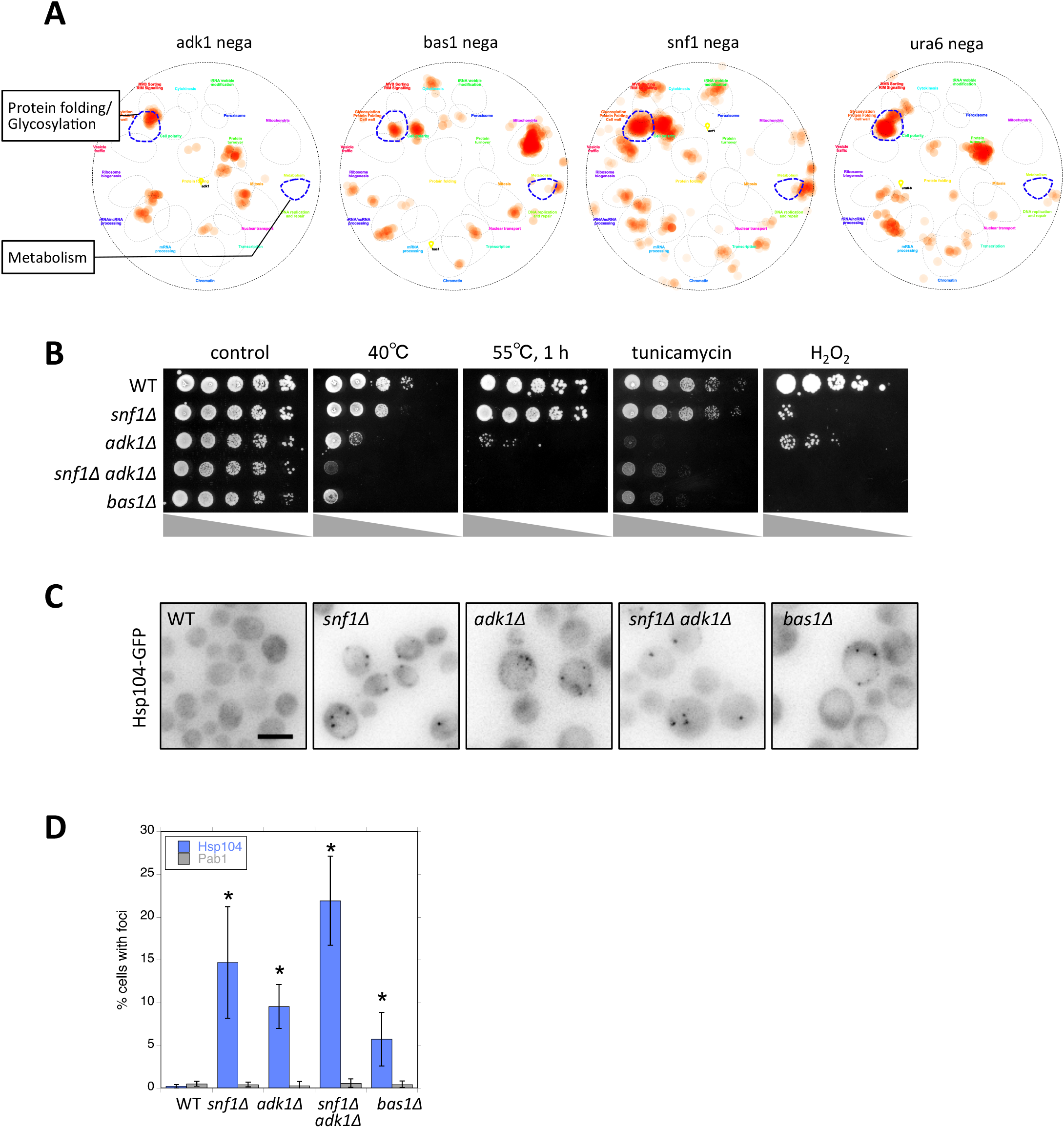
ATP homeostasis is required to prevent protein aggregation. (A) Functional landscape of known interactors of ATP mutants. Negative genetic interactors of the indicated gene were derived from the SGD database (https://www.yeastgenome.org/) (Cherry, Hong et al., 2012) and overlaid on a functional map based on the global genetic interaction network of the yeast genome (Baryshnikova, 2016, Usaj et al., 2017). *URA6* encodes an uridylate kinase that is essential for viability, which also exhibits adenylate kinase activity. (B) Each strain of the indicated genotype was serially diluted (five-fold), spotted on SC + 2% glucose medium, and grown under the indicated stress conditions. Photos were taken after 2-3 days. (C) Formation of Hsp104-GFP foci in ATP homeostasis mutants. The GFP signal (inverted grayscale) was imaged in the log phase culture of the indicated mutant cells expressing Hsp104-GFP at 35°C. (D) Quantification of data shown in (C). Data from similar experiments using strains expressing Pab1-GFP, instead of Hsp104-GFP, were also plotted. Values are the mean ± 1SD (error bars). Asterisks indicate a significant difference from WT (*p* < 0.05) (N=3–4). White scale bar = 5 µm.

To examine possible defects in protein folding and turnover (*i*.*e*., proteostasis), we challenged these mutants with various proteotoxic stresses. We found negligible growth defects in ATP mutants under normal growth conditions with 2% glucose at 30°C (control in Fig. 3B), suggesting that a high level of ATP is not necessary for cellular growth. However, the *adk1* and *bas1* mutants both exhibited severe growth defects with a high temperature of 40°C, 1 hour of heat shock at 55°C, or in the presence of 0.5 µg/ml of the glycosylation inhibitor tunicamycin or 2 mM H_2_O_2_. The *SNF1* deletion increased the stress sensitivity of *adk1Δ* (Fig.3B). This sensitivity to proteotoxic stress suggests that ATP homeostasis mutants are defective in some aspects of proteostasis. We found that all four mutants tested contained significantly increased numbers of Hsp104-GFP foci, a marker of protein aggregation (Josefson, Andersson et al., 2017) (Fig. 3C, D). In contrast to Hsp104-GFP foci, Pab1-GFP, a marker of stress granule (SG) assembly (Hoyle, Castelli et al., 2007), did not form foci in ATP mutants, suggesting that protein aggregation and SG assembly are regulated in a distinct manner (Fig. 3D). These analyses identified abnormal protein aggregation as a common defect associated with ATP homeostasis mutants for the first time.

### The transient depletion of ATP leads to the formation of protein aggregates

To examine whether ATP depletion triggers protein aggregation in living yeast, we artificially depleted cellular ATP levels by replacing glucose with 2-deoxyglucose (2DG), a strong inhibitor of glycolysis, in media and monitored protein aggregation using Hsp104-GFP as a marker of protein aggregation (Josefson et al., 2017) in wild-type cells. We previously showed that ATP levels were almost completely depleted 2 minutes after the 2DG treatment (Takaine et al., 2019), which was also confirmed biochemically (Fig. S3B). Within 15 min of the 2DG treatment, more than 20% of cells contained Hsp104-GFP foci indicative of protein aggregation (Fig. 4A, B). These protein aggregations were retained for hours after refeeding of glucose (Fig. 4A, B), suggesting that the dissolution kinetics of Hsp104-GFP were significantly slow. This contrasts intracellular ATP, which recovers to normal levels within 1 min of glucose refeeding (Takaine et al., 2019).

**Fig 4.**
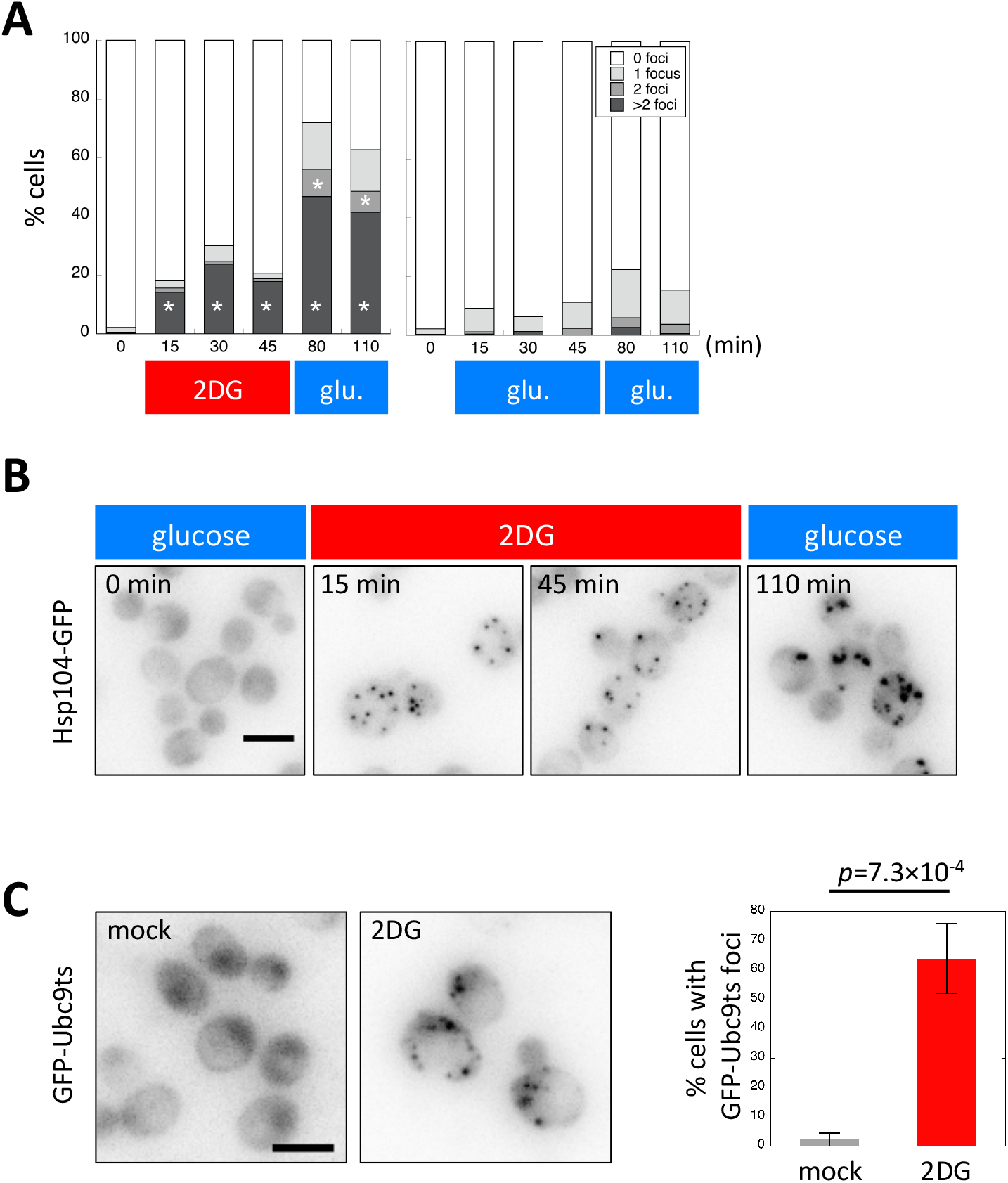
ATP depletion triggers protein aggregation in living yeast cells. (A) The formation of Hsp104-GFP foci after ATP depletion. Wild-type cells expressing Hsp104-GFP were grown to the log phase at 35°C in medium containing 2% glucose. At the time point of 0 min, medium was replaced with 20 mM 2DG (red) or 2% glucose (as a control; blue). Cells were released back to media containing 2% glucose at the time point of 50 min. Cells were imaged at the indicated time points, classified, and scored according to the number of Hsp104-GFP foci. Values are the mean (N = 3). Asterisks indicate a significant difference from the 2% glucose treatment (*p* < 0.05). (B) Representative images of cells analyzed in (A). (C) Formation of Ubc9-ts foci after ATP depletion. Cells expressing GFP-Ubc9-ts under an inducible *GAL* promoter were grown in medium containing 2% galactose (SC-gal) at 33°C, and medium was then exchanged with 2DG or SC-gal. After 30 min, cells were imaged and scored for the number of GFP-Ubc9-ts foci. Representative images (inverted grayscale) are shown on the left and summarized on the right. Values are the mean ± 1SD (error bars) (N = 4). White scale bar = 5 µm.

To further confirm whether a high level of ATP is required for protein solubility, we also tested the Ubc9-ts protein, a model protein that is prone to aggregation (Kaganovich, Kopito et al., 2008), and found that ATP depletion by the 2DG treatment triggered Ubc9-ts protein aggregation (Fig. 4C). Therefore, not only Hsp104-GFP-positive intrinsic proteins, but also extrinsic model proteins aggregate after ATP depletion.

SG are assembled in budding yeast cells under stress conditions, such as glucose depletion (Hoyle et al., 2007). In contrast to the formation of Hsp104-GFP foci, ATP depletion after the 2DG treatment did not instantly trigger the formation of SG (Fig. S8). Consistent with recent findings (Jain, Wheeler et al., 2016), the present results suggest that SG formation requires ATP. We also noted that Hsp104-GFP foci and SG did not co-localize, indicating that these structures are derived from distinct mechanisms (Fig. S8). Therefore, the artificial depletion of ATP may trigger abnormal protein aggregation in living yeast cells.

### ATP homeostasis is required for the protection of cells from cytotoxicity caused by protein aggregation

Protein aggregation is often associated with neurodegenerative diseases, such as Alzheimer’s, Huntington’s, and Parkinson’s diseases (Eftekharzadeh, Hyman et al., 2016). Mitochondrial failure has also been associated with many neurodegenerative diseases; however, it currently remains unclear whether energy failure causes protein aggregation because mitochondria also produce cytotoxic reactive oxygen species (ROS). (Bhat, Dar et al., 2015, Pathak, Berthet et al., 2013). The abnormal aggregation of α-synuclein has been implicated in Parkinson’s disease (Lashuel, Overk et al., 2013). To clarify whether ATP prevents the formation of cytotoxic protein aggregation, we examined the toxicity of α-synuclein-GFP (Syn-GFP) in budding yeast. As reported previously, the expression of Syn-GFP exhibited negligible toxicity against wild-type yeast when expressed under the inducible *GAL1* promotor (Fig. 5A) (Outeiro & Lindquist, 2003, Sharma, Brandis et al., 2006, Wijayanti, Watanabe et al., 2015). However, *snf1Δ, adk1Δ*, and *bas1Δ* were hypersensitive to the expression of Syn-GFP (Fig. 5A). We also found that *rpn4Δ*, which encodes a key transcription factor for proteasomal subunits (Xie & Varshavsky, 2001), was very sensitive to Syn-GFP (Fig. 5A), which is consistent with the concept that Syn-GFP is degraded in the ubiquitin-proteasomal pathway in yeast (Tofaris, Kim et al., 2011, Wijayanti et al., 2015). We then visualized the cellular localization of Syn-GFP. Consistent with previous findings (Willingham, Outeiro et al., 2003), Syn-GFP expressed in yeast mainly localized to the plasma membrane (Fig. 5B, C). In addition to the plasma membrane, we found that Syn-GFP localized to punctate structures, most likely corresponding to protein aggregation (Fig. 5B, C). Punctate structures were not as obvious in *rpn4Δ* cells defective in proteasomes, suggesting that the accumulation of Syn-GFP puncta was not simply due to defective degradation.

**Fig 5.**
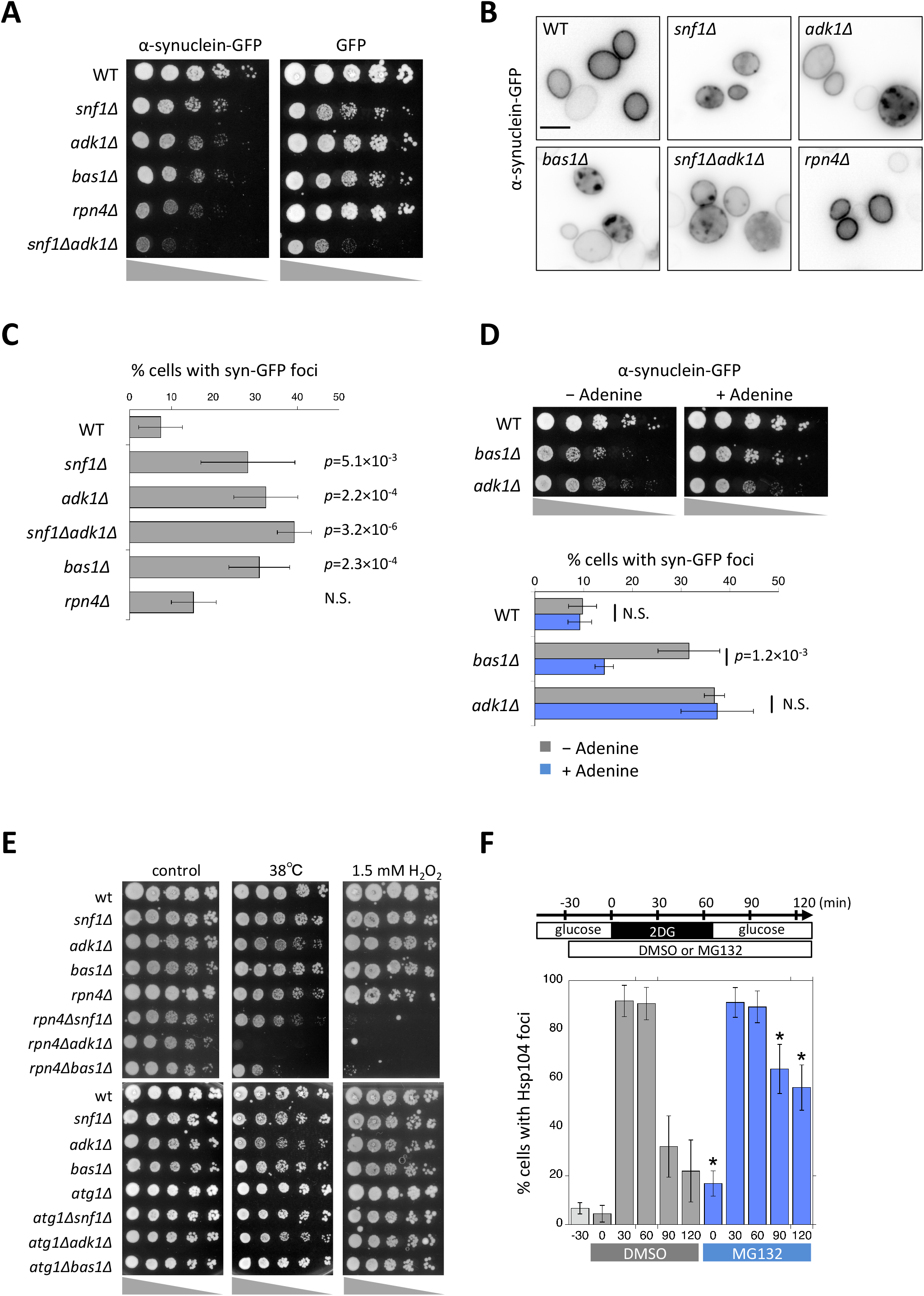
Aggregation and cytotoxicity of α-synuclein depends on ATP homeostasis. (A) Each strain of the indicated genotype was transformed with an expression vector carrying α-synuclein-GFP or GFP, serially diluted (five-fold), spotted on SC + 2% galactose plates, and then grown at 30°C for 3 days. (B) The localization of α-synuclein-GFP in ATP mutants. Cells were grown on galactose plates at 30°C for more than 42 h and then imaged. Representative images of α-synuclein-GFP (inverted grayscale) are shown. (C) Quantification of the data shown in (B). Cells were classified and scored for the localization pattern of α-synuclein-GFP. The percentage of cells showing α-synuclein-GFP foci are plotted. Data are the mean ± 1SD (error bars) from 3–6 independent observations. N = 33–380 cells were scored in each measurement. *P* values versus WT are shown. N.S., no significance (*p* value > 0.05). (D) *(top)* Each strain of the indicated genotype was transformed with an expression vector carrying α-synuclein-GFP and grown on galactose plates containing 0 mM (− Adenine) or 0.3 mM (+ Adenine) adenine at 30°C for 3 days. *(bottom)* Cells were grown on galactose plates in the absence or presence of adenine at 30°C for 41-45 h and then imaged. The percentage of cells showing α-synuclein-GFP foci was plotted. Data are the mean ± 1SD (error bars) from 5 independent transformants. N = 53–258 cells were scored in each measurement. *P* values versus “– Adenine” are shown. (E) Each strain of the indicated genotype was serially diluted (five-fold), spotted on SC + 2% glucose medium, and grown under the indicated stress conditions. (F) Cells of the drug-sensitive strain Y13206 were grown to the log phase at 37°C in medium containing 2% glucose and supplemented with 0.1% DMSO or 0.1% DMSO plus 42 µM MG132 at *t* = –30 min. At *t* = 0 min, these cells were washed and released in medium containing 20 mM 2DG ± MG132, and cells were then washed and released again in medium containing 2% glucose ± MG132. Cells were imaged at the indicated time points and scored for the number of Hsp104-GFP foci. Data are the mean ± 1SD (error bars). Asterisks indicate a significant difference from DMSO (*p* < 0.02) (N=3).

To investigate whether a high level of ATP protects cells from toxic protein aggregation, we added extra adenine to the medium (Fig. 5D). A previous study demonstrated that the addition of 300 µM adenine to the medium increased ATP levels from 4 to 5.5 mM in wild-type cells and from 1 to 4 mM in *bas1Δ* cells (Gauthier et al., 2008) (similar results are shown in Fig.1E), but induced negligible or no changes in *adk1Δ* cells (from 2 to 2 mM) (Gauthier et al., 2008). Consistent with our hypothesis, the addition of adenine reduced Syn-GFP toxicity and aggregation in *bas1Δ*, but not *adk1Δ* cells (Fig. 5D). Thus, a high level of ATP prevented Syn-GFP aggregation and toxicity.

We examined another model protein involved in neurodegenerative diseases. PolyQ containing the huntingtin protein is susceptible to aggregation and has been implicated in Huntington’s disease (Jiang, Poirier et al., 2005). We investigated the toxicity of Htt103Q, a mutant form of the huntingtin protein that is also susceptible to aggregation and causes cytotoxicity in yeast (Meriin, Zhang et al., 2002). Consistent with the concept that a high level of ATP prevents protein aggregation, the ATP homeostasis mutants *snf1Δ, adk1Δ, snf1Δ adk1Δ*, and *bas1Δ* were very sensitive to Htt103Q expression (Fig. S9).

### Proteasomes are essential for the removal of protein aggregates induced by ATP depletion

Protein aggregation caused by ATP depletion was cytotoxic (Figs. 3B and 5A) and was not easily dissolved after ATP repletion (Fig. 4). To identify a pathway that is essential for the removal of aggregates, we examined the involvement of proteasomes and autophagy.

The deletion of *RPN4*, which encodes a transcription factor of proteasomal genes (Xie & Varshavsky, 2001), down-regulated proteasomal activity (Kruegel, Robison et al., 2011) and resulted in synthetic growth defects with *adk1Δ, snf1Δ, bas1Δ* at a high temperature of 38°C and in the presence of H_2_O_2_ (Fig. 5E). In contrast to proteasomes, autophagy did not appear to have genetic interactions with the above mutants (Fig. 5E). The deletion of an essential component of the autophagic pathway, *ATG1* did not affect the sensitivity of *adk1Δ, snf1Δ, bas1Δ* to a high temperature of 38°C or to H_2_O_2_ (Fig. 5E). We also did not observe the accumulation of Hsp104-GFP foci in the autophagy mutants *atg1Δ, atg8Δ*, and *atg13Δ* (not shown).

To investigate the involvement of proteasomes in the removal of protein aggregates after the transient depletion of ATP, we pretreated cells with the proteasomal inhibitor MG132 or DMSO and examined the kinetics for the formation of Hsp104-GFP foci after the 2DG treatment (Fig. 5F) using the drug-sensitive yeast strain Y13206 (Piotrowski, Li et al., 2017). Under both conditions, more than 90% of cells exhibited Hsp104-GFP foci within 30 min of the 2DG treatment. More than two-thirds of Hsp104-GFP foci dissolved in the DMSO control, while less than one-third dissolved in MG132-treated samples, indicating that proteasomes are required for the dissolution process (Fig. 5F).

## DISCUSSION

In the present study, we demonstrated for the first time that the Snf1 complex, budding yeast AMPK, is required for the stable maintenance of cellular ATP levels (ATP homeostasis) in collaboration with Adk1 (Fig. 6A). This function of the Snf1 complex in ATP homeostasis is independent of glucose levels in the medium and Mig1, the major transcriptional repressor involved in glucose repression (Fig. S1); therefore, this is distinct from its well-characterized role in adaptation to glucose limitations. The activity of the Snf1 kinase complex may be sharply tuned depending on the intracellular levels of adenine nucleotides or other metabolites indicative of cellular energy to prevent a rapid ATP catastrophe (Fig. 2), even in the presence of sufficient amounts of glucose. It is important to note that the reductions observed in intracellular ATP levels in *snf1Δ* cells in the presence of glucose were overlooked in previous biochemical analyses, again demonstrating the usefulness of QUEEN-based ATP imaging.

**Fig 6.**
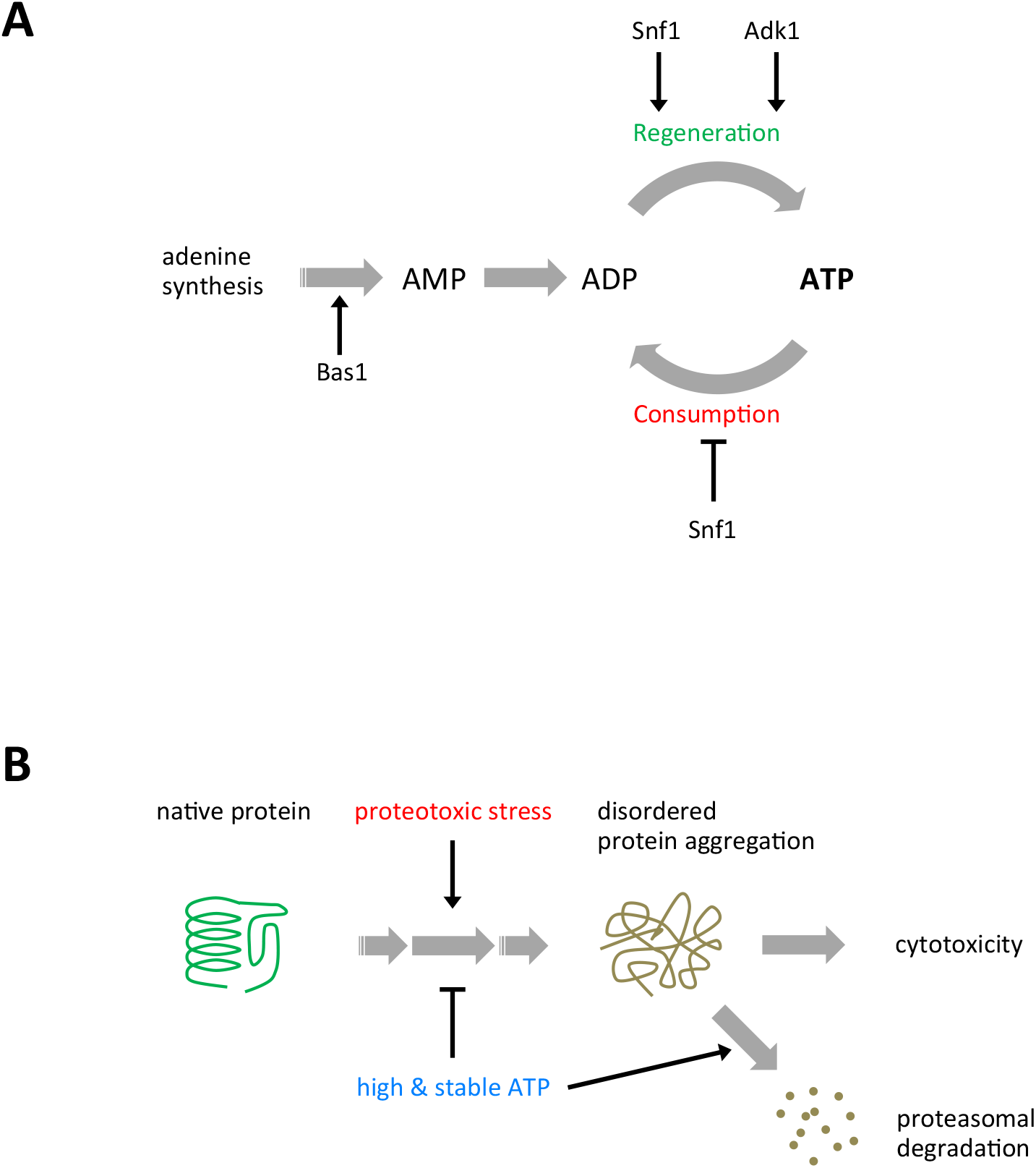
Models for ATP homeostasis and its role in proteostasis. (A) Schematic summary of the roles of Snf1, Adk1, and Bas1 in ATP homeostasis. (B) A schematic model for ATP homeostasis preventing cytotoxic protein aggregation.

Since the deletion of *BAS1* induced the greatest reduction in ATP levels and is epistatic to *snf1Δ*, a large pool size of adenine nucleotides is a prerequisite for ATP homeostasis. This assumption is reasonable because the pool size of recyclable ATP restricts ATP levels based on the rapid turnover rate of ATP. Bas1 maintains the pool size of ATP by balancing ATP synthesis and irreversible decreases, such as incorporation into RNA and DNA (following conversion to deoxy-ATP), degradation, and excretion in rapidly proliferating yeasts.

The decreases observed in ATP levels in the ATP mutant cells were confirmed by our biochemical measurements (Figs. 1 and S3). The biochemical assay also revealed that ADP levels decreased in ATP mutants, similar to ATP levels, and, as a consequence, ATP/ADP ratios, indicators of cellular energy charges, remained largely unchanged. On the other hand, the sum of ATP and ADP levels, indicators of the pool size of adenine nucleotides, decreased in the mutants, which may explain the instability observed in ATP levels. While cytosolic ATP levels were reduced and unstable in *snf1Δadk1Δ* cells, high levels of ATP may accumulate in intracellular membrane compartments (*e*.*g*., vacuoles, lysosomes, and mitochondria) that were not visualized by the QUEEN, which would increase the average ATP level of whole cells (Fig. 1B, C).

We also showed that key regulators of ATP homeostasis play roles in preventing cytotoxic protein aggregation in budding yeast (Fig. 6B). The common feature associated with these mutants is reduced ATP levels, suggesting that high ATP levels are essential for protein solubilization.

A proteomic study suggested that the main role of ATP changes depending on its level. At levels lower than 0.5 mM, ATP mainly serves as a substrate for enzymes, such as protein kinases and heat shock protein chaperones, whereas at levels higher than 2 mM, ATP may exert solubilizing effects on disordered proteins (Sridharan et al., 2019). ATP homeostasis may also be required to constantly drive proteasomal protein degradation, which requires high levels of ATP (Benaroudj, Zwickl et al., 2003, Tanaka, Waxman et al., 1983).

Previous biochemical measurements indicated that although ATP levels were lower in *adk1* and *bas1* mutants than in the wild type, these mutants still had ATP levels that were higher than 2 mM (Gauthier et al., 2008). This does not directly explain the accumulation of protein aggregates in these mutants (Fig. 3) because most proteins are expected to be soluble at > 2 mM ATP (Sridharan et al., 2019). In this study, we explained this discrepancy by using the biosensor-based ATP imaging technique we developed (Takaine et al., 2019, Yaginuma et al., 2014). The visualization of ATP dynamics in living cells at the single cell level revealed that ATP repeatedly undergoes transient depletion (the ATP catastrophe) in AMPK and ADK mutants, which explains how aggregation-prone proteins aggregate and cause cytotoxicity. A similar, but distinct, instability in ATP was observed in *bas1Δ*, further confirming that the ATP catastrophe causes protein aggregation. Although severe ATP depletion in these mutants was gradually recovered by as yet uncharacterized negative feedback regulation, the duration period of ∼15 min with reduced ATP levels may induce some proteins to form aggregates that last for generations.

A recent study revealed that forced energy (ATP) depletion from budding yeast cells induced extensive cytoplasmic reorganization, including increases in macromolecular (ribosome) crowding, the emergence of numerous membrane-less organelles, and the polymerization of eukaryotic translation initiation factor 2B (Marini et al., 2020). Another recent study showed that the depletion of ATP triggered the “viscoadaptation” of yeast cytoplasm by an as yet unknown mechanism, and induced increases in viscosity and decreases in biomolecular solubility, thereby enhancing phase separation (Persson et al., 2020). These findings imply that malfunctional ATP homeostasis may induce undesirable and dysregulated biomolecular assemblies driven by enhanced liquid-liquid phase separation and molecular crowding. The present study using ATP mutant cells is the first to demonstrate the consequence of failed ATP homeostasis under physiologically relevant conditions and highlights the biological significance of ATP homeostasis.

Many neurodegenerative diseases, such as Alzheimer’s, Huntington’s, and Parkinson’s diseases, are associated with protein aggregation (Eisele, Monteiro et al., 2015, Josefson et al., 2017). On the other hand, based on a large body of evidence, mitochondrial dysfunction and accompanying energy failure in nerve cells may result in many types of neurodegenerative diseases (Haelterman, Yoon et al., 2014, Pathak et al., 2013). Previous studies demonstrated that ATP levels in the brain were decreased in patients with early Parkinson’s disease (Mochel, N’Guyen et al., 2012) and also that ATP synthesis in the brain was not properly regulated in patients with early Huntington’s disease (Hattingen, Magerkurth et al., 2009) and in the corresponding mouse model (Mochel, Durant et al., 2012). Therefore, protein aggregation induced by the ATP catastrophe, as revealed in the present study, may link energy failure and protein aggregation, providing a comprehensive insight into the onset of neurodegenerative diseases. Further studies are warranted to clarify whether the ATP catastrophe also occurs in the neurons of patients at risk of neurodegenerative diseases or in the elderly. However, neither biochemical measurements nor mass spectrometry is capable of detecting the ATP catastrophe because of their insufficient time and space resolution. Therefore, an ATP imaging approach using the yeast system will be the leading model for elucidating the molecular mechanisms underlying ATP homeostasis and ATP catastrophe-induced protein aggregation.

A recent study reported that the activation of AMPK by metformin ameliorated the progression of Huntington’s disease in a mouse model (Arnoux, Willam et al., 2018), and the potential therapeutic use of metformin for neurodegenerative diseases is being discussed (Rotermund, Machetanz et al., 2018). Furthermore, the involvement of ATP and ADK in preventing the manifestation of Parkinson’s disease in mouse models and patients has been proposed (Garcia-Esparcia, Hernandez-Ortega et al., 2015, Nakano, Imamura et al., 2017). Protein aggregation induced by the ATP catastrophe may be a general mechanism for the development of proteinopathies. The present study using ATP imaging revealed a physiological consequence of a failure in ATP homeostasis in living cells for the first time and suggests that ATP homeostasis has potential as a target for preventing/treating neurodegenerative diseases.

## MATERIALS AND METHODS

### Yeast strains and plasmids

The budding yeast strains and plasmids used in the present study are listed in Supplementary Tables S1 and S2, respectively. These strains were constructed by a PCR-based method (Janke, Magiera et al., 2004) and genetic crosses. The yeast knockout strain collection was originally purchased from GE Healthcare (cat. # YSC1053).

### Media and cell culture

The standard technique for the yeast culture and manipulation was used (Guthrie & Fink, 2002). Synthetic medium (SC) was prepared as described by Hanscho et al. (Hanscho, Ruckerbauer et al., 2012). 2-Deoxy-D-glucose (2DG), tunicamycin, and MG132 were purchased from FUJIFILM Wako (cat. # 046-06483, 202-08241, and 139-18451, respectively). Tunicamycin and MG132 were dissolved in dimethylsulfoxide (DMSO) to make stock solutions (5 mg/ml and 42 mM, respectively). Cells were grown to the mid-log phase at 30°C in SC before imaging unless otherwise noted.

### Microscopy

Cells expressing Hsp104-GFP or GFP-Ubc9ts were concentrated by centrifugation and sandwiched between a slide and coverslip (No. 1.5 thickness, Matsunami, Osaka, Japan). Immobilized cells were imaged using an inverted fluorescent microscope (Eclipse Ti-E, Nikon) equipped with an Apo TIRF 100× Oil DIC N2/NA 1.49 objective lens and electron-multiplying charge-coupled device camera (iXon3 DU897E-CS0-#BV80, Andor) at approximately 25°C. The Hsp104-GFP and GFP-Ubc9ts fluorescent signal was collected from stacks of 11 *z*-sections spaced by 0.5 µm, and the maximum projections of the optical sections were shown in Figs. 4, 5, and S6. Cells expressing QUEEN were immobilized on a concanavalin A-coated 35-mm glass-bottomed dish (#3971-035, No. 1.5 thickness, IWAKI). The dish was filled with an excess amount of medium (4.5–5 ml) against the cell volume to minimize changes in the chemical compositions of the medium during observations. QUEEN fluorescence was acquired as previously described (Takaine et al., 2019). Cells expressing Syn-GFP were immobilized on a slide glass as described above, and the fluorescence signal was collected from a single *z*-plane using an inverted fluorescent microscope (Eclipse Ti2-E, Nikon, Tokyo, Japan) equipped with a CFI Plan Apoλ100× Oil DIC/NA1.45 objective lens and CMOS image sensor (DS-Qi2, Nikon). Images of cells were acquired from several fields of view for each experimental condition.

### Biochemical measurements of ATP and ADP

Whole cell extracts were prepared according to Seo et al. (Seo, Lau et al., 2017) with slight modifications. Mid-log cells were harvested in a 1.6-ml microtube by centrifugation and resuspended in 1 ml fresh SC or 40 mM 2DG medium (for ATP depletion). After a 10-min incubation at 30°C, a small fraction of cells (50 µl) was removed for the measurement of cell numbers and optical density, and the remaining cells were pelleted and resuspended in 0.75 ml of 90% acetone. The suspension was incubated at 90°C for 15 min to evaporate acetone. The remaining solution (30–35 µl) was centrifuged at 20,000×*g* at 4°C for 3 min. The supernatant was mixed with 450 µl of TE (10 mM Tris-HCl, pH 8.0, and 1 mM ethylenediaminetetraacetic acid). These extracts were stored at −80°C until analyzed. ATP and ADP levels were measured using the ATP determination kit (Invitrogen) and EnzyLight ADP assay kit (EADP-100, Funakoshi), respectively, according to the manufacturers’ instructions. Luminescence was measured using an Enspire multimode plate reader (PerkinElmer). All samples were assayed in duplicate. ATP and ADP levels were normalized for an optical density at 600 nm of the initial cell suspension assessed by BioSpectrometer (Eppendorf).

### Data analysis

Numerical data were plotted using KaleidaGraph software ver. 4.5.1 (Synergy Software) and R studio software ver. 3.4.1 (R Core Team, 2017). Means, SDs, and *p* values were calculated using Excel software (Microsoft, WA, US). Significance between two sets of data was tested using the unpaired one-tailed Welch’s *t*-test unless otherwise noted, and was indicated by an asterisk or *p* value. Data were sometimes represented by a dot plot that shows distribution characteristics in extensive detail. The horizontal bar in the dot plot indicates the average of each population. All measurements were repeated at least twice to confirm reproducibility.

ATP levels in yeast cells were estimated using QUEEN-based ratiometric imaging, as previously described (Takaine, 2019), (Takaine et al., 2019). The QUEEN ratio is proportional to ATP levels and pseudo-colored to reflect its value throughout the present study. The mean QUEEN ratio inside of a cell represents the intracellular ATP level of the cell.

The autocorrelation functions (ACF) of oscillations in the QUEEN ratio were calculated using R studio software. The apparent period of oscillation was estimated from the positive second peak of the correlation coefficient, which was outside the 95% confidence interval and significant (*p* < 0.05), rejecting the assumption that there is no correlation.

## ACKNOWLEDGMENTS

We are grateful to the Yeast Genetic Resource Center, Y. Ohya, K. Ohashi, H. Takagi, D. Watanabe, and J. Frydman for providing the yeast strains and plasmids. We thank the members of the Yoshida/Takaine laboratories for their support. This work was supported by JSPS grants 16H04781 (S.Y. and M.T.), 15K18525 (M.T.), and 19K06654 (M.T.) and the Takeda Science Foundation (S.Y.). This work was also supported by the joint research program of the Institute for Molecular and Cellular Regulation, Gunma University, Japan.

## AUTHOR CONTRIBUTIONS

M.T. and S.Y. conceived and designed the project. M.T. conducted experiments and the data analysis. H.I. provided a key reagent and expertise. M.T. and S.Y wrote the manuscript with input from H.I.

## SUPPLEMENTARY INFORMATION

### Supplementary figure legends

**Fig. S1.**
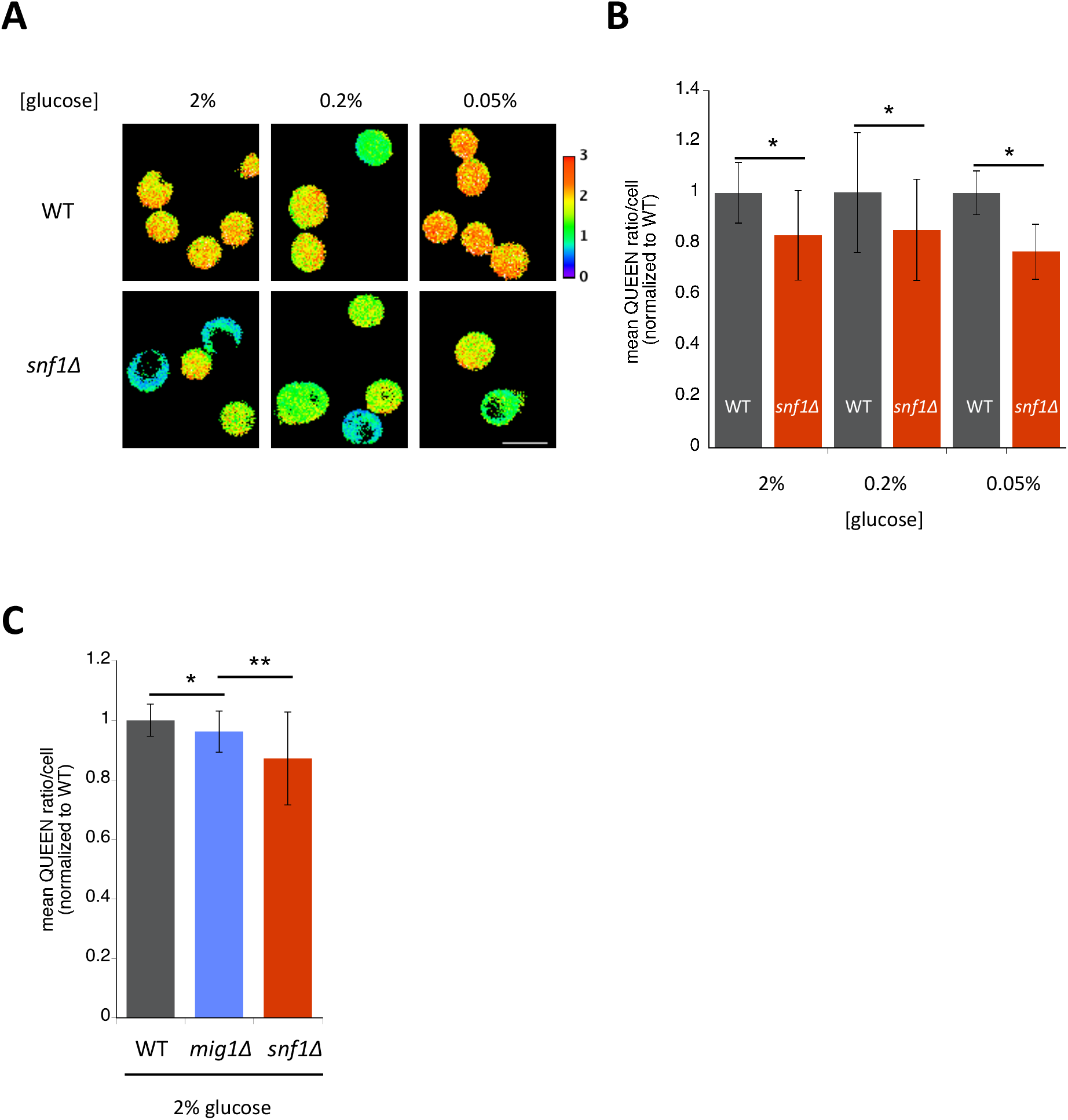
AMPK is involved in the maintenance of cellular ATP levels in non-starving cells. (A) Visualization of intracellular ATP levels in wild-type (WT) and *snf1Δ* cells using the ATP sensor QUEEN. Cells were grown to the mid-log phase in SC-H medium containing the indicated levels of glucose and then imaged. White scale bar = 5 µm. (B) *snf1Δ* cells had lower ATP levels than wild-type cells. The mean QUEEN ratio inside a single cell (mean QUEEN ratio/cell), which represents the intracellular ATP level of the cell, was quantified for each cell from the ratio image. Data are the mean of the cell population ± 1SD (error bar) normalized to wild-type cells. N = 105–193 cells were scored. Asterisks indicate a significant difference from WT (*p* < 0.05). (C) *mig1Δ* cells had slightly lower ATP levels than wild-type cells. Data are the mean of the cell population ± 1SD (error bar) normalized to wild-type cells. N = 190–231 cells were scored. Asterisks indicate a significant difference between the two strains.

**Fig. S2.**
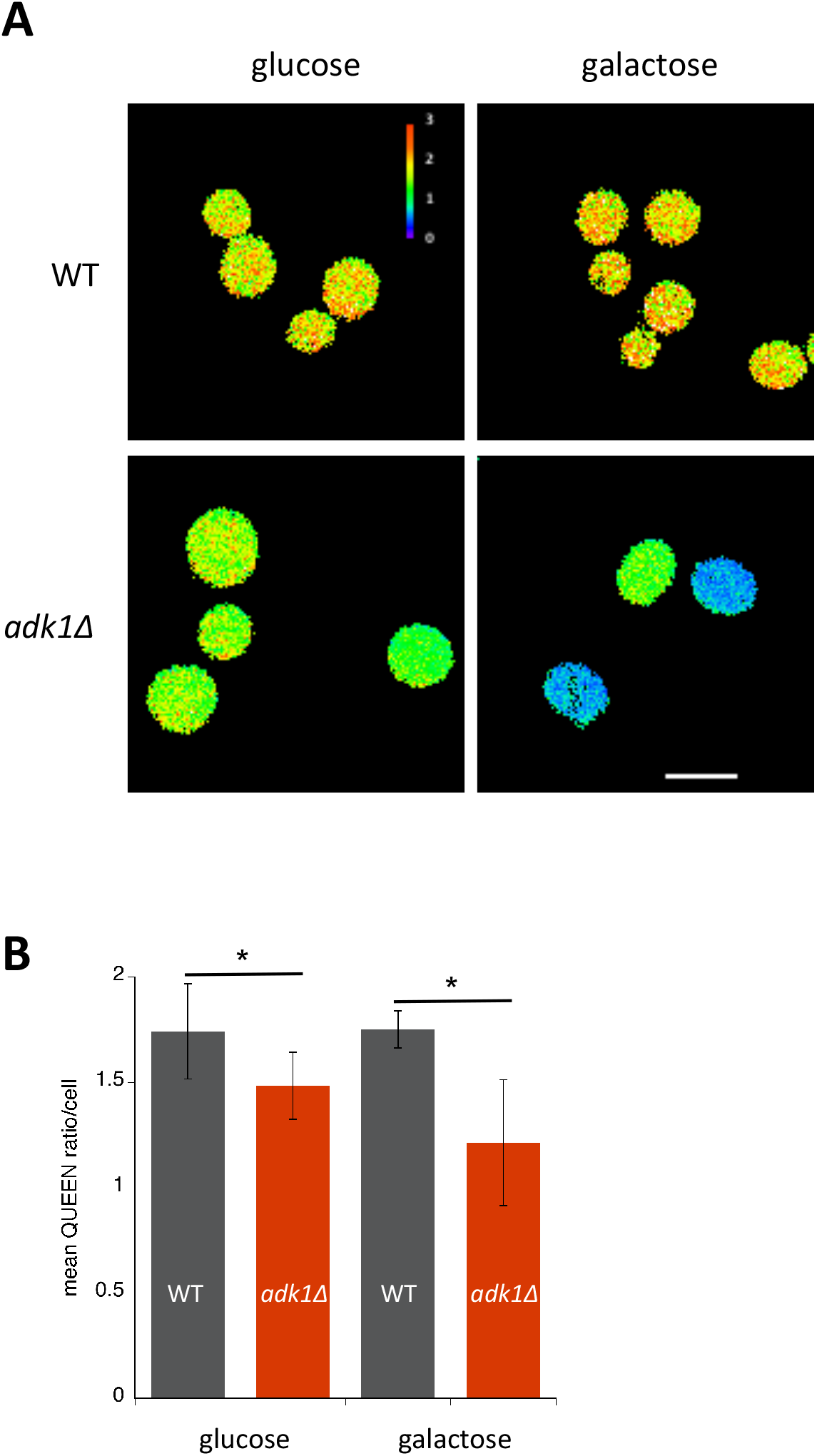
Adenylate kinase Adk1 is involved in the maintenance of cellular ATP levels. (A) QUEEN ratio images of wild-type and *adk1Δ* cells grown in 2% glucose or 2% galactose. White scale bar = 5 µm. (B) Quantification of data shown in (A). The mean QUEEN ratio/cell was quantified for each cell from ratio images. Values are the mean of the cell population ± 1SD (error bar). N = 182–236 cells were scored. Asterisks indicate a significant difference from WT (*p* < 0.05).

**Fig. S3.**
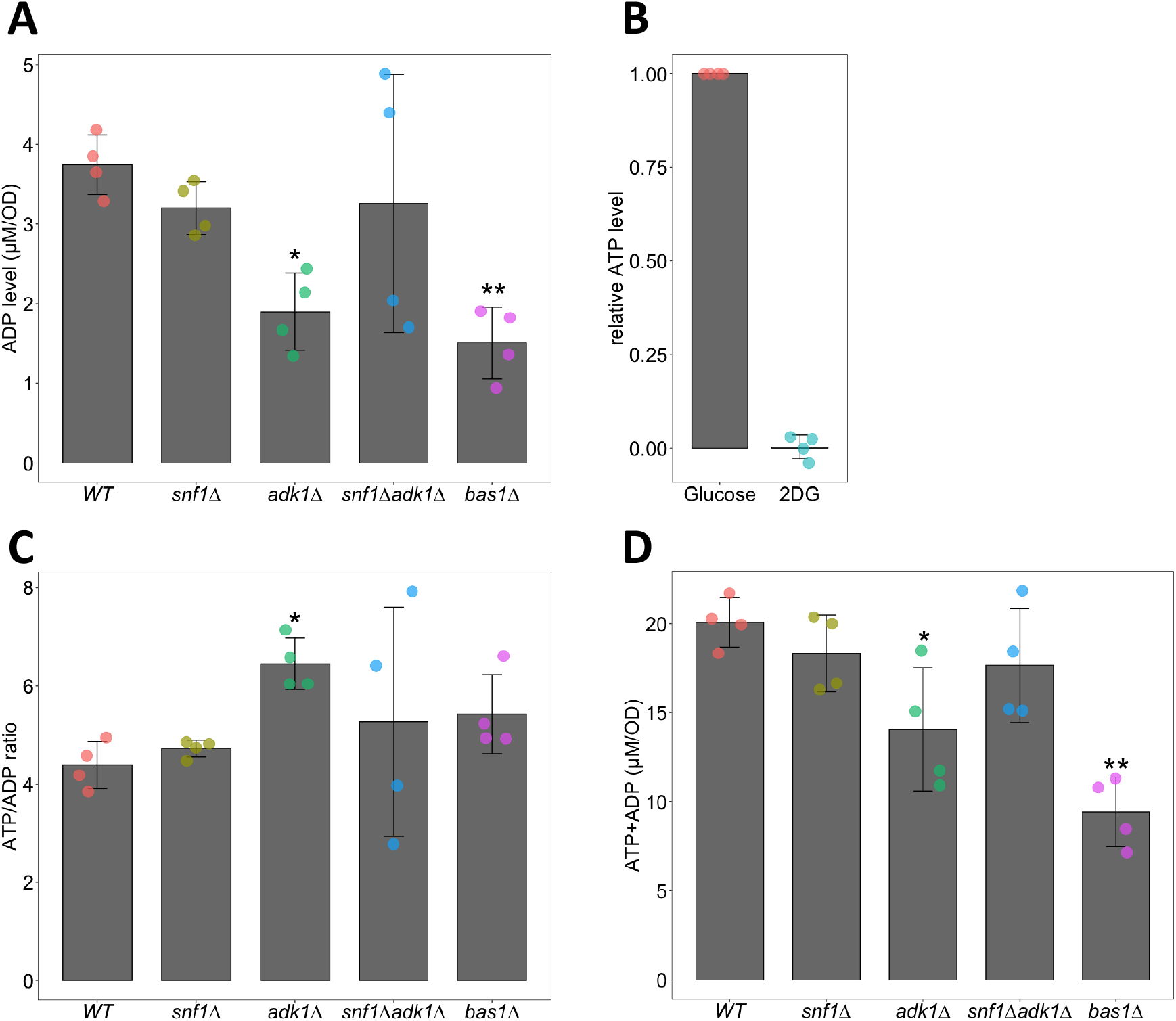
Biochemical measurements of ATP and ADP. (A) Biochemical measurements of cellular ADP levels. ADP levels in cells of the indicated genotypes were measured as described in the Materials and Methods. Data are the mean ± 1SD (error bars) (N = 4). Asterisks indicate *p* values versus WT: *=6.0×10^−4^, **=1.5×10^−4^. (B) 2DG depletes cellular ATP. ATP levels in WT cells incubated with medium containing 2% glucose or 40 mM 2DG for 10 min were biochemically measured. Relative ATP levels were expressed as the ratios of ATP levels in glucose-treated cells. Data are the mean ± 1SD (error bars) (N = 4). (C) ATP/ADP ratios in WT and ATP mutant cells. ATP/ADP ratios were calculated from ATP and ADP levels measured using biochemical assays. Data are the mean ± 1SD (error bars) (N = 4). An asterisk indicates a *p* value of 6.0×10^−4^ versus WT. (D) The sums of ATP and ADP levels in WT and ATP mutant cells were calculated from ATP and ADP levels measured using biochemical assays. Data are the mean ± 1SD (error bars) (N = 4). Asterisks indicate *p* values versus WT: *=1.6×10^−2^, **=1.0×10^−4^.

**Fig. S4.**
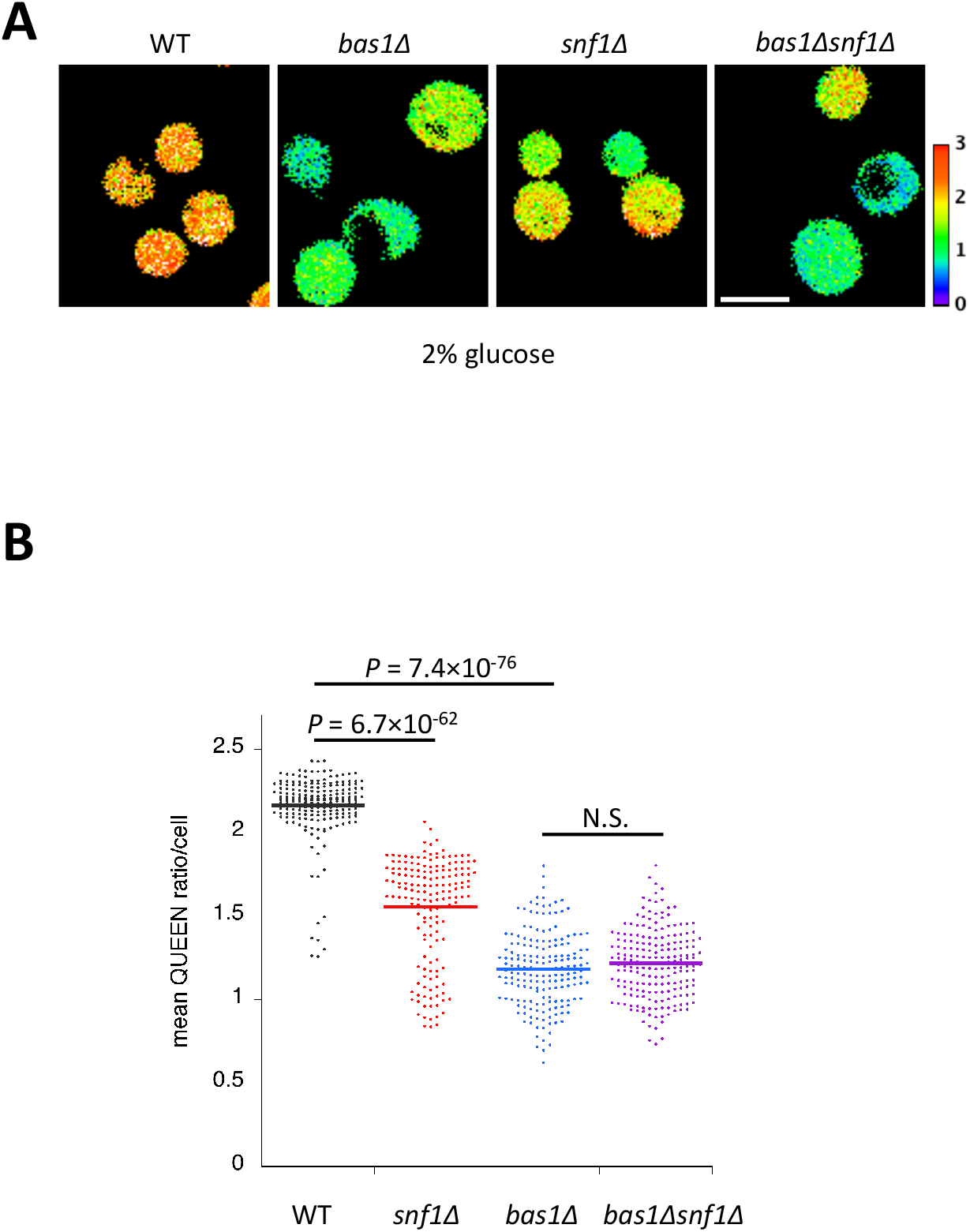
ATP levels in *bas1Δ snf1Δ* cells. (A) QUEEN ratio images of wild-type, *bas1Δ, snf1Δ*, and *bas1Δ snf1Δ* cells. (B) Quantification of data shown in (A). The mean QUEEN ratio/cell was quantified for each cell from ratio images. Data are shown as a dot plot. The horizontal bar in the plot indicates the mean of each population. Significance between two sets of data was tested using the unpaired two-tailed Welch’s *t*-test and indicated by *p* values. N.S., no significance (*p*-value > 0.05).

**Fig. S5.**
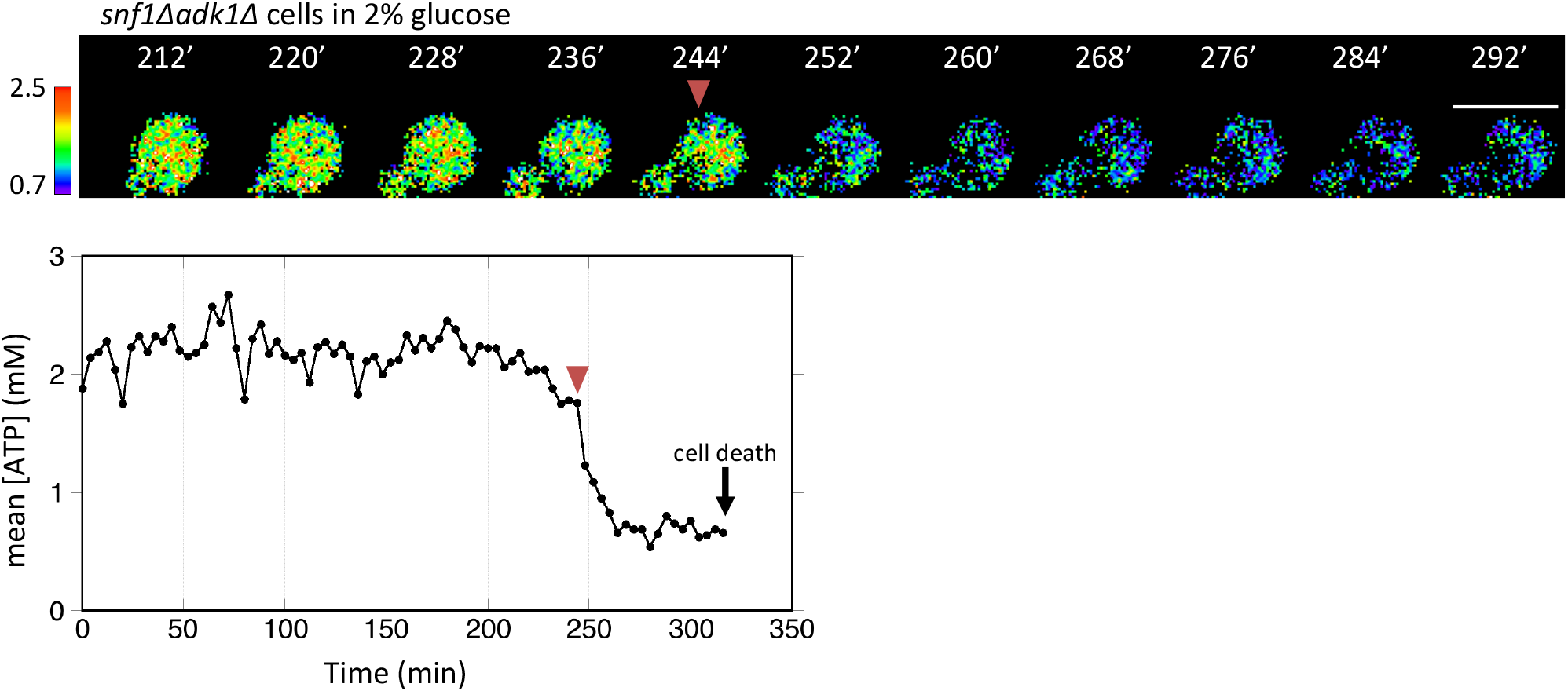
Time-lapse imaging of QUEEN in *snf1*Δ *adk1*Δ cells. Time-lapse imaging of QUEEN in *snf1*Δ *adk1*Δ cells in 2% glucose medium. An example of a *snf1*Δ *adk1*Δ cell showing an irreversible decrease in the QUEEN ratio (indicated by an arrowhead). In this case, the cell eventually died (indicated by an arrow). Images at the representative time points were shown. The ATP level was plotted at the bottom. White scale bar = 5 µm.

**Fig. S6.**
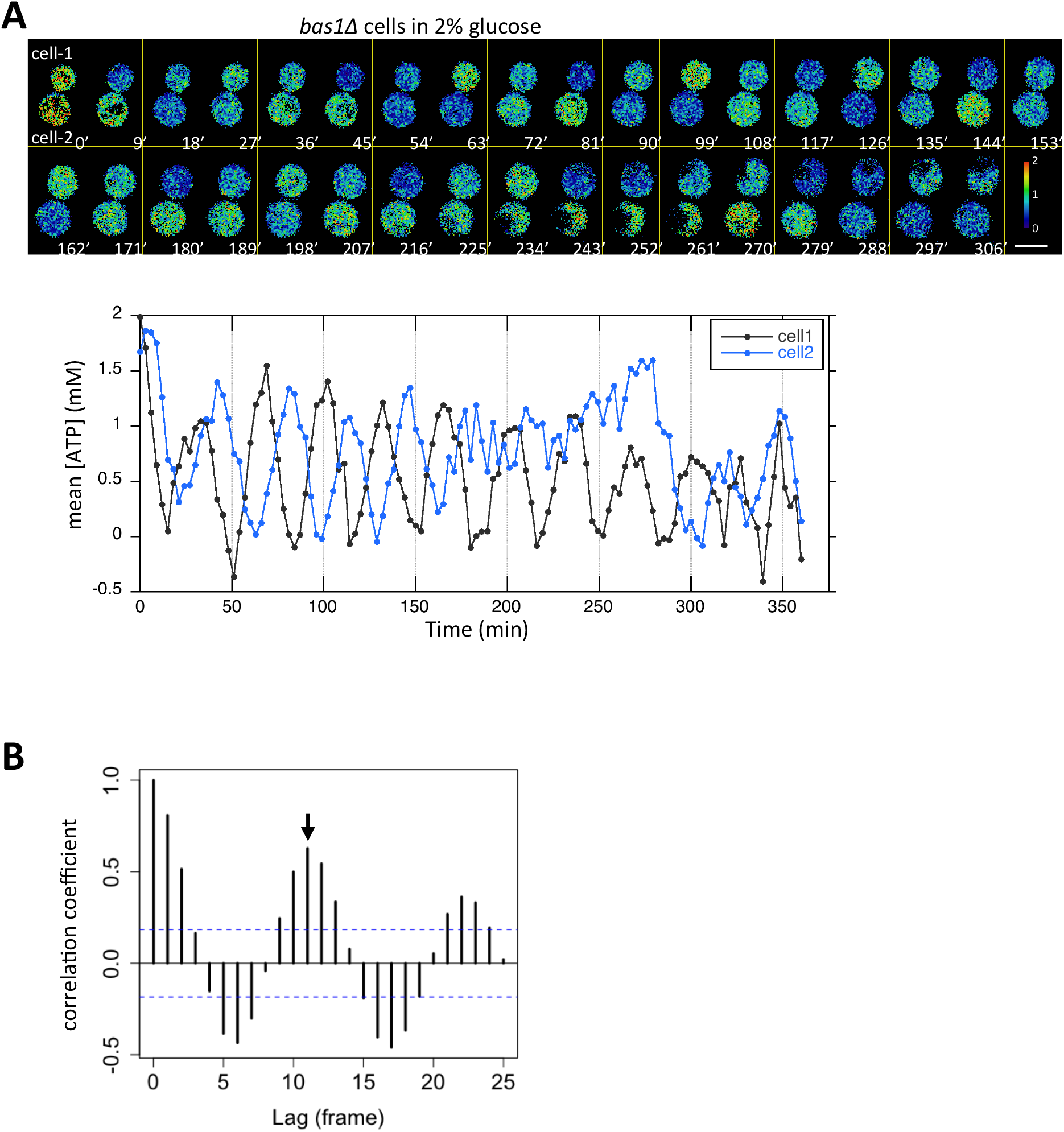
Oscillatory behavior of the ATP level visualized in *bas1Δ* cells. (A) Another example of *bas1Δ* cells showing an oscillating QUEEN ratio. In this case, cytokinesis had just been completed at t = 0 min. ATP levels in cell-1 and cell-2 were plotted at the bottom. See also Movie S4. White scale bar = 5 µm. (B) Autocorrelation function of the QUEEN ratio calculated from the data on cell-1 in Fig. 3E. Blue dotted lines indicate the 95% confidence interval. An arrow indicates the second peak of the correlation and corresponds to the apparent period.

**Fig. S7.**
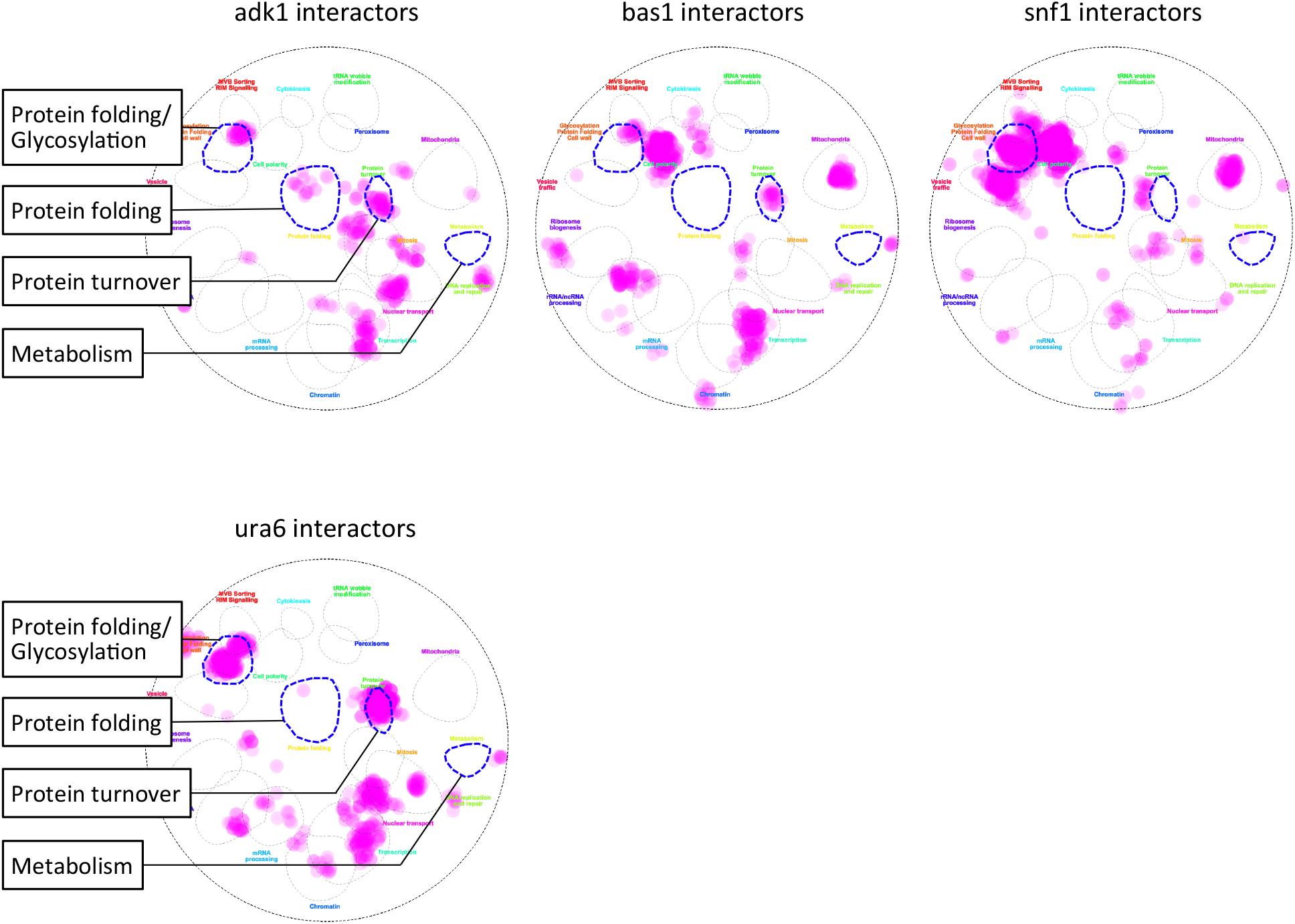
*In silico* analysis of interactors of ATP mutants. Genetic and physical interactors of the indicated genes were derived from the SGD database (https://www.yeastgenome.org/) (Cherry et al., 2012) and overlaid on a functional map based on the global genetic interaction network of the yeast genome (Baryshnikova, 2016, Usaj et al., 2017). *URA6* encodes an uridylate kinase essential for viability, which also exhibits adenylate kinase activity.

**Fig. S8.**
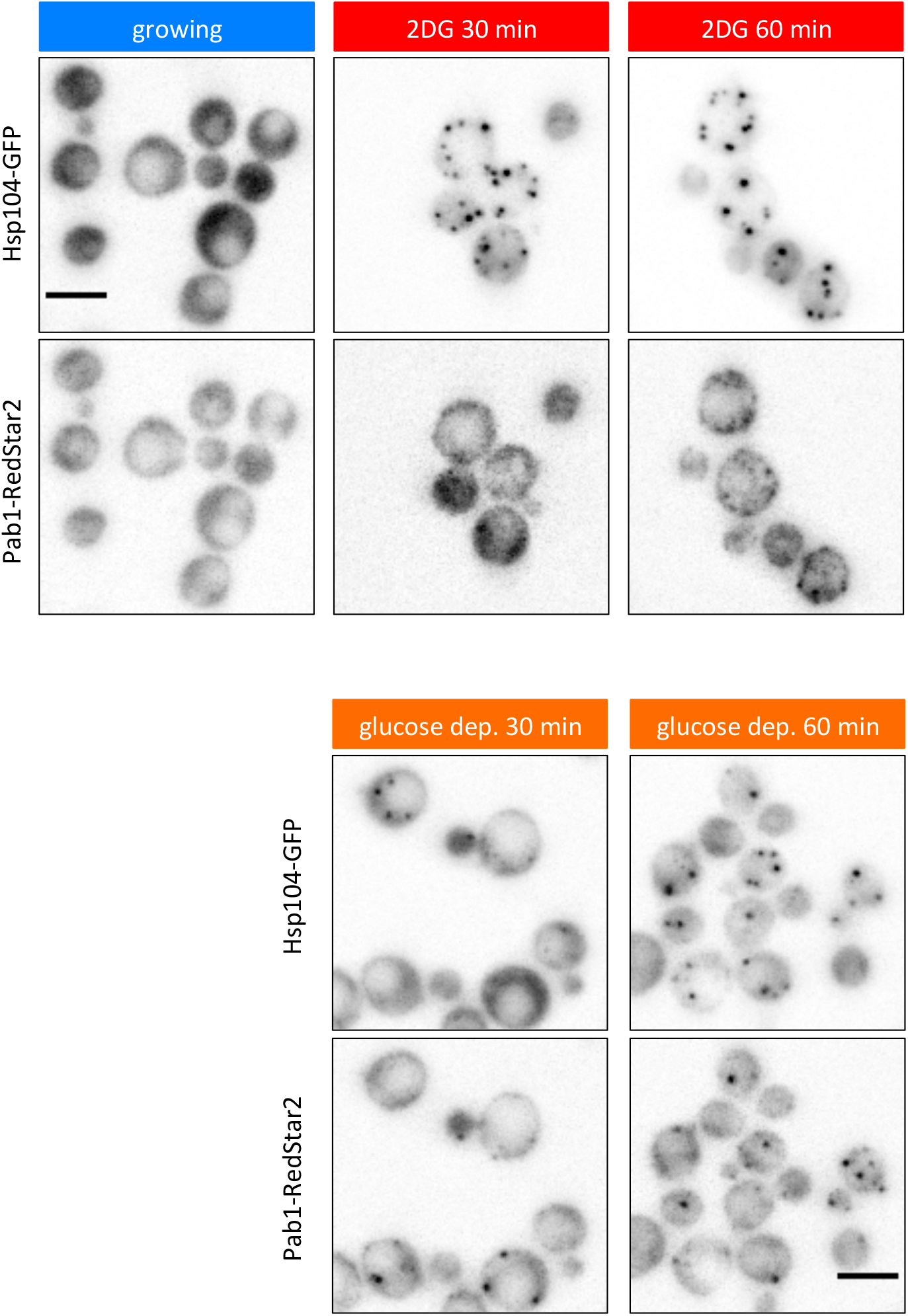
Simultaneous observation of Hsp104 and Pab1 foci. Wild-type cells expressing Hsp104-GFP and Pab1-RedStar2 were grown to the log phase at 37°C in medium containing 2% glucose. Cells were washed and released either in medium containing 20 mM 2DG (top) or in medium lacking glucose (bottom), and then imaged after 30 and 60 min. White scale bar = 5 µm.

**Fig. S9.**
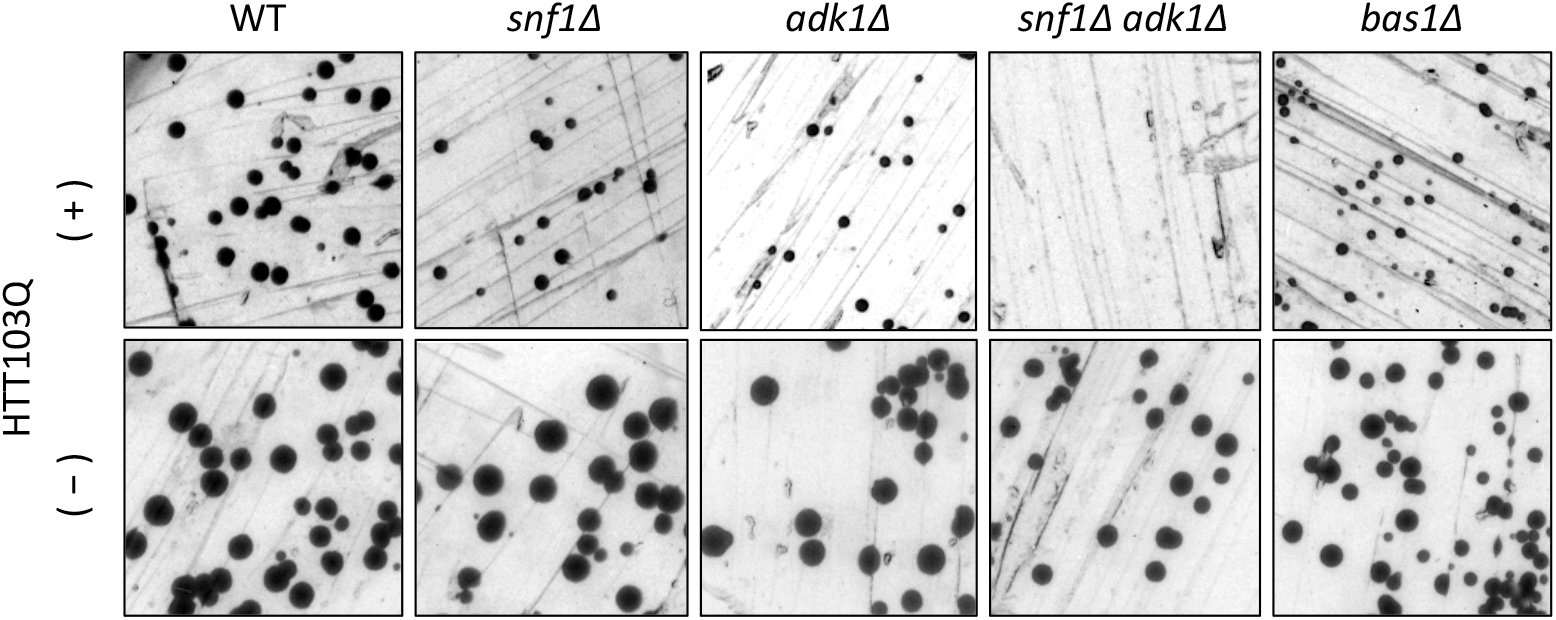
Cytotoxicity of polyQ containing the huntingtin protein in wild-type and ATP-mutant yeast cells. Each strain of the indicated genotype was transformed with an expression vector carrying Htt103Q and grown on glucose ((-), no induction) or raffinose ((+), leaky expression) plates at 30°C for 3 days.

**Table S1.** Strains used in the present study

| Name | Genotype | Source | Figure |
| --- | --- | --- | --- |
| MTY3008 | *snf1Δ::kanMX6 leu2Δ0 lys2Δ0 ura3Δ0* | Lab stock | 1C, 3B,<br>5A-C, 5E,<br>S9 |
| MTY3015 | *his3Δ1 leu2Δ0 lys2Δ0 ura3Δ0* | Lab stock | 1C, 1F,<br>3B, 5A-E,<br>S9 |
| MTY3049 | *adk1Δ::kanMX6 his3Δ1 leu2Δ0 lys2Δ0 ura3Δ0* | Lab stock | 1C, 3B,<br>5A-E, S9 |
| MTY3118 | *bas1Δ::kanMX6 leu2Δ0 lys2Δ0 ura3Δ0* | Lab stock | 1F, 3B,<br>5A-E, S9 |
| MTY3143 | *his3Δ1:: 2×pRS303-P<sub>TEF</sub>-QUEEN-2m-T<sub>CYC1</sub> leu2Δ0 lys2Δ0 ura3Δ0* | This study | S2 |
| MTY3149 | *adk1Δ::kanMX6 his3Δ1:: 2×pRS303-P<sub>TEF</sub>-QUEEN-2m-T<sub>CYC1</sub> leu2Δ0 lys2Δ0 ura3Δ0* | This study | S2 |
| MTY3264 | *his3Δ1:: 3×pRS303-P<sub>TEF</sub>-QUEEN-2m-T<sub>CYC1</sub> leu2Δ0 lys2Δ0 ura3Δ0*<br>*MYO1-3mCherry-hphMX6* | Takaine et al., 2019 | 1A-B, S1,<br>S4 |
| MTY3270 | *bas1Δ::kanMX6 his3Δ1:: 3×pRS303-P<sub>TEF</sub>-QUEEN-2m-T<sub>CYC1</sub> leu2Δ0 lys2Δ0 ura3Δ0*<br>*MYO1-3mCherry-hphMX6* | This study | 1D, 1E,<br>2E, S4, S6 |
| MTY3293 | *adk1Δ::kanMX6 his3Δ1:: 3×pRS303-P<sub>TEF</sub>-QUEEN-2m-T<sub>CYC1</sub> leu2Δ0 lys2Δ0 ura3Δ0*<br>*MYO1-3mCherry-hphMX6* | This study | 1A-B |
| MTY3355 | *adk1Δ::kanMX6 snf1Δ::natNT2 his3Δ1:: 3×pRS303-P<sub>TEF</sub>-QUEEN-2m-T<sub>CYC1</sub> leu2Δ0 lys2Δ0 ura3Δ0*<br>*MYO1-3mCherry-hphMX6* | This study | 1A-B,<br>2A-D, S5 |
| MTY3371 | *snf1Δ::kanMX6 his3Δ1:: 3×pRS303-P<sub>TEF</sub>-QUEEN-2m-T<sub>CYC1</sub> leu2Δ0 lys2Δ0 ura3Δ0*<br>*MYO1-3mCherry-hphMX6* | This study | 1A-B, S1,<br>S4 |
| MTY3412 | *adk1Δ::kanMX6 snf1Δ::natNT2 leu2Δ0 lys2Δ0 ura3Δ0* | This study | 1C, 3B,<br>5A-E, S9 |
| MTY3417 | *mig1Δ::natNT2 his3Δ1:: 3×pRS303-P<sub>TEF</sub>-QUEEN-2m-T<sub>CYC1</sub> leu2Δ0 lys2Δ0 ura3Δ0*<br>*MYO1-3mCherry-hphMX6* | This study | S1C |
| MTY3420 | *Hsp104-yeGFP-hphNT1 his3Δ1 leu2Δ0 lys2Δ0 ura3Δ0* | This study | 3C, D, 4A,<br>B |
| MTY3421 | *snf1Δ::kanMX6 Hsp104-yeGFP-hphNT1 his3Δ1 leu2Δ0 lys2Δ0 ura3Δ0* | This study | 3C, D |
| MTY3422 | *bas1Δ::kanMX6 Hsp104-yeGFP-hphNT1 his3Δ1 leu2Δ0 lys2Δ0 ura3Δ0* | This study | 3C, D |
| MTY3424 | *adk1Δ::kanMX6 Hsp104-yeGFP-hphNT1 his3Δ1 leu2Δ0 lys2Δ0 ura3Δ0* | This study | 3C-D |
| MTY3425 | *adk1Δ::kanMX6 snf1Δ::natNT2 Hsp104-yeGFP-hphNT1 his3Δ1 leu2Δ0 lys2Δ0 ura3Δ0* | This study | 3C, D |
| MTY3489 | *atg1Δ::kanMX6 his3Δ1 leu2Δ0 lys2Δ0 ura3Δ0* | Lab stock | 5E |
| MTY3493 | *snq2Δ::KILeu2; pdr3Δ::KLura3; pdr1Δ::natMX4; can1Δ::STE2prSp_his5 lyp1Δ his3Δ1 leu2Δ0 ura3Δ0 met15Δ0 LYS2+* | From Y. Ohya, (Piotrowski et al., 2017) | Parental strain of MTY3501 |
| MTY3501 | *Hsp104-yeGFP-hphNT1 snq2Δ::KILeu2; pdr3Δ::KLura3; pdr1Δ::natMX4; can1Δ::STE2prSp_his5 lyp1Δ his3Δ1 leu2Δ0 ura3Δ0 met15Δ0 LYS2+* | This study | 5F |
| MTY3503 | *Pab1- yeGFP-hphNT1 his3Δ1 leu2Δ0 lys2Δ0 ura3Δ0* | This study | 3D |
| MTY3504 | *Hsp104-yeGFP-hphNT1 Pab1-RedStar2-natNT2 his3Δ1 leu2Δ0 lys2Δ0 ura3Δ0* | This study | S8 |
| MTY3505 | *snf1Δ::kanMX6 Pab1- yeGFP-hphNT1 his3Δ1 leu2Δ0 lys2Δ0 ura3Δ0* | This study | 3D |
| MTY3506 | *adk1Δ::kanMX6 Pab1- yeGFP-hphNT1 his3Δ1 leu2Δ0 lys2Δ0 ura3Δ0* | This study | 3D |
| MTY3507 | *adk1Δ::kanMX6 snf1Δ::natNT2 Pab1- yeGFP-hphNT1 his3Δ1 leu2Δ0 lys2Δ0 ura3Δ0* | This study | 3D |
| MTY3508 | *bas1Δ::kanMX6 Pab1- yeGFP-hphNT1 his3Δ1 leu2Δ0 lys2Δ0 ura3Δ0* | This study | 3D |
| MTY3513 | *atg1Δ::natNT2 adk1Δ::kanMX6 his3Δ1 leu2Δ0 lys2Δ0 ura3Δ0* | This study | 5E |
| MTY3514 | *atg1Δ::natNT2 snf1Δ::kanMX6 his3Δ1 leu2Δ0 lys2Δ0 ura3Δ0* | This study | 5E |
| MTY3515 | *atg1Δ::natNT2 bas1Δ::kanMX6 his3Δ1 leu2Δ0 lys2Δ0 ura3Δ0* | This study | 5E |
| MTY3516 | *rpn4Δ::natNT2 adk1Δ::kanMX6 his3Δ1 leu2Δ0 lys2Δ0 ura3Δ0* | This study | 5E |
| MTY3517 | *rpn4Δ::natNT2 snf1Δ::kanMX6 his3Δ1 leu2Δ0 lys2Δ0 ura3Δ0* | This study | 5E |
| MTY3518 | *rpn4Δ::natNT2 bas1Δ::kanMX6 his3Δ1 leu2Δ0 lys2Δ0 ura3Δ0* | This study | 5E |
| MTY3525 | *rpn4Δ::kanMX6 his3Δ1 leu2Δ0 lys2Δ0 ura3Δ0* | Lab stock | 5A-C, 5E |

**Table S2.** Plasmids used in the present study

| Name | Structure | Source | Purpose |
| --- | --- | --- | --- |
| MTP3091 | pESC-Leu-GFP-Ubc9-ts | Judith Frydman (pJF1089) | Fig. 4C |
| MTP3088 | p426 103Q GAL | Addgene (cat.# 1188) | Fig. S9 |
| MTP3108 | pYES2-α-synuclein-GFP | H. Takagi (Wijayanti et al., 2015) | Fig. 5A-D |
| MTP3090 | pYES2-GFP | K. Ohashi | Fig. 5A |

## Captions for supplementary movies

**Movie S1**

Time-lapse imaging of QUEEN in *snf1*Δ*adk1*Δ cells in 2% glucose medium. Corresponding to the data shown in Fig. 2A. The QUEEN ratio decreased twice (116 and 132 min) within a short interval. White scale bar = 5 µm.

**Movie S2**

Another example of *snf1*Δ*adk1*Δ cells showing a sudden decrease in the QUEEN ratio. Corresponding to the data shown in Fig. 2B. The QUEEN ratio decreased twice (180 and 356 min) with a long interval. White scale bar = 5 µm.

**Movie S3**

Oscillatory behavior of the QUEEN ratio in *bas1Δ* cells. Corresponding to the data shown in Fig. 2E. Left, the QUEEN ratio image; middle, Myo1-mCherry (inverted grayscale image); right, bright field image. Images were taken every 4 min. White scale bar = 5 µm.

**Movie S4**

Another example of *bas1Δ* cells showing an oscillating QUEEN ratio. Corresponding to the data shown in Supplementary Fig. S6A. Left, the QUEEN ratio image; right, bright field image. In this case, cytokinesis had already been completed at t = 0 min. Images were taken every 3 min. White scale bar = 5 µm.

## References

Arnoux I, Willam M, Griesche N, Krummeich J, Watari H, Offermann N, Weber S, Narayan Dey P, Chen C, Monteiro O, Buettner S, Meyer K, Bano D, Radyushkin K, Langston R, Lambert JJ, Wanker E, Methner A, Krauss S, Schweiger S et al. (2018) Metformin reverses early cortical network dysfunction and behavior changes in Huntington’s disease. Elife 7

Baryshnikova A (2016) Systematic Functional Annotation and Visualization of Biological Networks. Cell Syst 2: 412–21

Benaroudj N, Zwickl P, Seemuller E, Baumeister W, Goldberg AL (2003) ATP hydrolysis by the proteasome regulatory complex PAN serves multiple functions in protein degradation. Mol Cell 11: 69–78

Bhat AH, Dar KB, Anees S, Zargar MA, Masood A, Sofi MA, Ganie SA (2015) Oxidative stress, mitochondrial dysfunction and neurodegenerative diseases; a mechanistic insight. Biomed Pharmacother 74: 101–10

Carlson M (1999) Glucose repression in yeast. Curr Opin Microbiol 2: 202–7

Cherry JM, Hong EL, Amundsen C, Balakrishnan R, Binkley G, Chan ET, Christie KR, Costanzo MC, Dwight SS, Engel SR, Fisk DG, Hirschman JE, Hitz BC, Karra K, Krieger CJ, Miyasato SR, Nash RS, Park J, Skrzypek MS, Simison M et al. (2012) Saccharomyces Genome Database: the genomics resource of budding yeast. Nucleic Acids Res 40: D700–5

Daignan-Fornier B, Fink GR (1992) Coregulation of purine and histidine biosynthesis by the transcriptional activators BAS1 and BAS2. Proc Natl Acad Sci U S A 89: 6746–50

Denis V, Boucherie H, Monribot C, Daignan-Fornier B (1998) Role of the myb-like protein bas1p in Saccharomyces cerevisiae: a proteome analysis. Mol Microbiol 30: 557–66

Edelman AM, Blumenthal DK, Krebs EG (1987) Protein serine/threonine kinases. Annu Rev Biochem 56: 567–613

Eftekharzadeh B, Hyman BT, Wegmann S (2016) Structural studies on the mechanism of protein aggregation in age related neurodegenerative diseases. Mech Ageing Dev 156: 1–13

Eisele YS, Monteiro C, Fearns C, Encalada SE, Wiseman RL, Powers ET, Kelly JW (2015) Targeting protein aggregation for the treatment of degenerative diseases. Nat Rev Drug Discov 14: 759–80

Garcia-Esparcia P, Hernandez-Ortega K, Ansoleaga B, Carmona M, Ferrer I (2015) Purine metabolism gene deregulation in Parkinson’s disease. Neuropathol Appl Neurobiol 41: 926–40

Gauthier S, Coulpier F, Jourdren L, Merle M, Beck S, Konrad M, Daignan-Fornier B, Pinson B (2008) Co-regulation of yeast purine and phosphate pathways in response to adenylic nucleotide variations. Mol Microbiol 68: 1583–94

Ghillebert R, Swinnen E, Wen J, Vandesteene L, Ramon M, Norga K, Rolland F, Winderickx J (2011) The AMPK/SNF1/SnRK1 fuel gauge and energy regulator: structure, function and regulation. FEBS J 278: 3978–90

Guthrie C, Fink GR (2002) Guide to yeast genetics and molecular and cell Biology: Part C. Gulf Professional Publishing,

Haelterman NA, Yoon WH, Sandoval H, Jaiswal M, Shulman JM, Bellen HJ (2014) A Mitocentric View of Parkinson’s Disease. Annual Review of Neuroscience 37: 137–159

Hanscho M, Ruckerbauer DE, Chauhan N, Hofbauer HF, Krahulec S, Nidetzky B, Kohlwein SD, Zanghellini J, Natter K (2012) Nutritional requirements of the BY series of Saccharomyces cerevisiae strains for optimum growth. FEMS Yeast Res 12: 796–808

Hardie DG, Carling D, Carlson M (1998) The AMP-activated/SNF1 protein kinase subfamily: metabolic sensors of the eukaryotic cell? Annu Rev Biochem 67: 821–55

Hardie DG, Schaffer BE, Brunet A (2016) AMPK: An Energy-Sensing Pathway with Multiple Inputs and Outputs. Trends Cell Biol 26: 190–201

Hattingen E, Magerkurth J, Pilatus U, Mozer A, Seifried C, Steinmetz H, Zanella F, Hilker R (2009) Phosphorus and proton magnetic resonance spectroscopy demonstrates mitochondrial dysfunction in early and advanced Parkinson’s disease. Brain 132: 3285–97

Hayes MH, Peuchen EH, Dovichi NJ, Weeks DL (2018) Dual roles for ATP in the regulation of phase separated protein aggregates in Xenopus oocyte nucleoli. Elife 7: e35224

Hedbacker K, Carlson M (2008) SNF1/AMPK pathways in yeast. Front Biosci 13: 2408–20

Herzig S, Shaw RJ (2017) AMPK: guardian of metabolism and mitochondrial homeostasis. Nature Reviews Molecular Cell Biology 19: 121

Hoyle NP, Castelli LM, Campbell SG, Holmes LE, Ashe MP (2007) Stress-dependent relocalization of translationally primed mRNPs to cytoplasmic granules that are kinetically and spatially distinct from P-bodies. J Cell Biol 179: 65–74

Jain S, Wheeler JR, Walters RW, Agrawal A, Barsic A, Parker R (2016) ATPase-Modulated Stress Granules Contain a Diverse Proteome and Substructure. Cell 164: 487–98

Janke C, Magiera MM, Rathfelder N, Taxis C, Reber S, Maekawa H, Moreno-Borchart A, Doenges G, Schwob E, Schiebel E, Knop M (2004) A versatile toolbox for PCR-based tagging of yeast genes: new fluorescent proteins, more markers and promoter substitution cassettes. Yeast 21: 947–62

Janssen E, Dzeja PP, Oerlemans F, Simonetti AW, Heerschap A, de Haan A, Rush PS, Terjung RR, Wieringa B, Terzic A (2000) Adenylate kinase 1 gene deletion disrupts muscle energetic economy despite metabolic rearrangement. EMBO J 19: 6371–81

Jiang H, Poirier MA, Ross CA (2005) A structure-based analysis of huntingtin mutant polyglutamine aggregation and toxicity: evidence for a compact beta-sheet structure. Human Molecular Genetics 14: 765–774

Josefson R, Andersson R, Nyström T (2017) How and why do toxic conformers of aberrant proteins accumulate during ageing? Essays In Biochemistry 61: 317

Kaganovich D, Kopito R, Frydman J (2008) Misfolded proteins partition between two distinct quality control compartments. Nature 454: 1088–95

Kruegel U, Robison B, Dange T, Kahlert G, Delaney JR, Kotireddy S, Tsuchiya M, Tsuchiyama S, Murakami CJ, Schleit J, Sutphin G, Carr D, Tar K, Dittmar G, Kaeberlein M, Kennedy BK, Schmidt M (2011) Elevated proteasome capacity extends replicative lifespan in Saccharomyces cerevisiae. PLoS Genet 7: e1002253

Lashuel HA, Overk CR, Oueslati A, Masliah E (2013) The many faces of alpha-synuclein: from structure and toxicity to therapeutic target. Nat Rev Neurosci 14: 38–48

Ljungdahl PO, Daignan-Fornier B (2012) Regulation of amino acid, nucleotide, and phosphate metabolism in Saccharomyces cerevisiae. Genetics 190: 885–929

Marini G, Nuske E, Leng W, Alberti S, Pigino G (2020) Reorganization of budding yeast cytoplasm upon energy depletion. Mol Biol Cell 31: 1232–1245

Meriin AB, Zhang X, He X, Newnam GP, Chernoff YO, Sherman MY (2002) Huntington toxicity in yeast model depends on polyglutamine aggregation mediated by a prion-like protein Rnq1. J Cell Biol 157: 997–1004

Mochel F, Durant B, Meng X, O’Callaghan J, Yu H, Brouillet E, Wheeler VC, Humbert S, Schiffmann R, Durr A (2012) Early alterations of brain cellular energy homeostasis in Huntington disease models. J Biol Chem 287: 1361–70

Mochel F, N’Guyen TM, Deelchand D, Rinaldi D, Valabregue R, Wary C, Carlier PG, Durr A, Henry PG (2012) Abnormal response to cortical activation in early stages of Huntington disease. Mov Disord 27: 907–10

Nakano M, Imamura H, Sasaoka N, Yamamoto M, Uemura N, Shudo T, Fuchigami T, Takahashi R, Kakizuka A (2017) ATP Maintenance via Two Types of ATP Regulators Mitigates Pathological Phenotypes in Mouse Models of Parkinson’s Disease. EBioMedicine 22: 225–241

Outeiro TF, Lindquist S (2003) Yeast cells provide insight into alpha-synuclein biology and pathobiology. Science 302: 1772–5

Patel A, Malinovska L, Saha S, Wang J, Alberti S, Krishnan Y, Hyman AA (2017) ATP as a biological hydrotrope. Science 356: 753–756

Pathak D, Berthet A, Nakamura K (2013) Energy failure: does it contribute to neurodegeneration? Ann Neurol 74: 506–16

Persson LB, Ambati VS, Brandman O (2020) Cellular Control of Viscosity Counters Changes in Temperature and Energy Availability. Cell

Piotrowski JS, Li SC, Deshpande R, Simpkins SW, Nelson J, Yashiroda Y, Barber JM, Safizadeh H, Wilson E, Okada H, Gebre AA, Kubo K, Torres NP, LeBlanc MA, Andrusiak K, Okamoto R, Yoshimura M DeRango-Adem E, van Leeuwen J, Shirahige K et al. (2017) Functional annotation of chemical libraries across diverse biological processes. Nature Chemical Biology 13: 982

Pu Y, Li Y, Jin X, Tian T, Ma Q, Zhao Z, Lin SY, Chen Z, Li B, Yao G, Leake MC, Lo CJ, Bai F (2019) ATP-Dependent Dynamic Protein Aggregation Regulates Bacterial Dormancy Depth Critical for Antibiotic Tolerance. Mol Cell 73: 143–156 e4

R Core Team (2017) R: A Language and Environment for Statistical Computing. In Vienna, Austria: R Foundation for Statistical Computing

Rotermund C, Machetanz G, Fitzgerald JC (2018) The Therapeutic Potential of Metformin in Neurodegenerative Diseases. Front Endocrinol (Lausanne) 9: 400

Seo AY, Lau P-W, Feliciano D, Sengupta P, Gros MAL, Cinquin B, Larabell CA, Lippincott-Schwartz J (2017) AMPK and vacuole-associated Atg14p orchestrate µ-lipophagy for energy production and long-term survival under glucose starvation. eLife 6: e21690

Sharma N, Brandis KA, Herrera SK, Johnson BE, Vaidya T, Shrestha R, DebBurman SK (2006) α-synuclein budding yeast model. Journal of Molecular Neuroscience 28: 161–178

Sridharan S, Kurzawa N, Werner T, Gunthner I, Helm D, Huber W, Bantscheff M, Savitski MM (2019) Proteome-wide solubility and thermal stability profiling reveals distinct regulatory roles for ATP. Nat Commun 10: 1155

Takaine M (2019) QUEEN-based Spatiotemporal ATP Imaging in Budding and Fission Yeast. Bio-Protocol 9: e3320

Takaine M, Ueno M, Kitamura K, Imamura H, Yoshida S (2019) Reliable imaging of ATP in living budding and fission yeast. J Cell Sci 132

Tanaka K, Waxman L, Goldberg AL (1983) ATP serves two distinct roles in protein degradation in reticulocytes, one requiring and one independent of ubiquitin. J Cell Biol 96: 1580–5

Tofaris GK, Kim HT, Hourez R, Jung JW, Kim KP, Goldberg AL (2011) Ubiquitin ligase Nedd4 promotes alpha-synuclein degradation by the endosomal-lysosomal pathway. Proc Natl Acad Sci U S A 108: 17004–9

Usaj M, Tan Y, Wang W, VanderSluis B, Zou A, Myers CL, Costanzo M, Andrews B, Boone C (2017) TheCellMap.org: A Web-Accessible Database for Visualizing and Mining the Global Yeast Genetic Interaction Network. G3 (Bethesda) 7: 1539–1549

Wijayanti I, Watanabe D, Oshiro S, Takagi H (2015) Isolation and functional analysis of yeast ubiquitin ligase Rsp5 variants that alleviate the toxicity of human α-synuclein. The Journal of Biochemistry 157: 251–260

Willingham S, Outeiro TF, DeVit MJ, Lindquist SL, Muchowski PJ (2003) Yeast genes that enhance the toxicity of a mutant huntingtin fragment or alpha-synuclein. Science 302: 1769–72

Wilson WA, Hawley SA, Hardie DG (1996) Glucose repression/derepression in budding yeast: SNF1 protein kinase is activated by phosphorylation under derepressing conditions, and this correlates with a high AMP:ATP ratio. Curr Biol 6: 1426–34

Xiao B, Heath R, Saiu P, Leiper FC, Leone P, Jing C, Walker PA, Haire L, Eccleston JF, Davis CT, Martin SR, Carling D, Gamblin SJ (2007) Structural basis for AMP binding to mammalian AMP-activated protein kinase. Nature 449: 496–500

Xie Y, Varshavsky A (2001) RPN4 is a ligand, substrate, and transcriptional regulator of the 26S proteasome: a negative feedback circuit. Proc Natl Acad Sci U S A 98: 3056–61

Yaginuma H, Kawai S, Tabata KV, Tomiyama K, Kakizuka A, Komatsuzaki T, Noji H, Imamura H (2014) Diversity in ATP concentrations in a single bacterial cell population revealed by quantitative single-cell imaging. Sci Rep 4: 6522

